# Ability of nucleoside-modified mRNA to encode HIV-1 envelope trimer nanoparticles

**DOI:** 10.1101/2021.08.09.455714

**Authors:** Zekun Mu, Kevin Wiehe, Kevin O. Saunders, Rory Henderson, Derek W. Cain, Robert Parks, Diana Martik, Katayoun Mansouri, Robert J. Edwards, Amanda Newman, Xiaozhi Lu, Shi-Mao Xia, Mattia Bonsignori, David Montefiori, Qifeng Han, Sravani Venkatayogi, Tyler Evangelous, Yunfei Wang, Wes Rountree, Ying Tam, Christopher Barbosa, S. Munir Alam, Wilton B. Williams, Norbert Pardi, Drew Weissman, Barton F. Haynes

## Abstract

The success of nucleoside-modified mRNAs in lipid nanoparticles (mRNA-LNP) as COVID-19 vaccines heralded a new era of vaccine development. For HIV-1, multivalent envelope (Env) trimer protein nanoparticles are superior immunogens compared to trimers alone for priming of broadly neutralizing antibody (bnAb) B cell lineages. The successful expression of complex multivalent nanoparticle immunogens with mRNAs has not been demonstrated. Here we show that mRNAs can encode antigenic Env trimers on ferritin nanoparticles that initiate bnAb precursor B cell expansion and induce serum autologous tier 2 neutralizing activity in bnAb precursor VH + VL knock-in mice. Next generation sequencing demonstrated acquisition of critical mutations, and monoclonal antibodies that neutralized heterologous HIV-1 isolates were isolated. Thus, mRNA-LNP can encode complex immunogens and are of use in design of germline-targeting and sequential boosting immunogens for HIV-1 vaccine development.

## INTRODUCTION

The recent success of nucleoside-modified mRNA COVID-19 vaccines encoding SARS-CoV- 2 trimeric spike protein has demonstrated the robust nature of the mRNA vaccine platform (Baden et al., 2020; Buschmann et al., 2021; Sahin et al., 2020). In addition to success with clinically-approved COVID-19 spike trimer vaccines, pre-clinical success has been demonstrated with nucleoside-modified mRNA encapsulated in lipid nanoparticles (mRNA-LNP) expression of Zika prM-E (Pardi et al., 2017), influenza hemagglutinin (Pardi et al., 2018a; Pardi et al., 2018c), and HIV-1 envelope (Env) in gp120 monomeric or gp140 trimeric forms (Mu et al., 2021; Pardi et al., 2018a; Saunders et al., 2021). However, recent studies have shown that protein trimer multimers presented on a nanoparticle (NP) scaffold may be advantageous as immunogens, particularly for engaging B cell receptors (BCRs) of HIV-1 broadly neutralizing antibody (bnAb) B cell precursors that are rare or have low affinity (Abbott et al., 2018; Havenar-Daughton et al., 2018; Kato et al., 2020; Saunders et al., 2019; Tokatlian et al., 2019).

HIV-1 bnAbs may be disfavored by the immune system due to their unusual characteristics of long heavy-chain complementarity-determining region 3 (HCDR3) loops and polyreactivity or autoreactivity that predispose bnAbs to immune tolerance control (Havenar-Daughton et al., 2018; Haynes et al., 2019; Haynes et al., 2005; Haynes et al., 2012; Haynes et al., 2016; Huang et al., 2020; Saunders et al., 2019; Steichen et al., 2019; Zhang et al., 2016). Thus, the biology of HIV- bnAbs has necessitated a strategy whereby the unmutated common ancestor (UCA) or germline (GL) precursor of bnAb B cell lineages is targeted with priming immunogens to expand the bnAb precursor pool (Haynes et al., 2019; Haynes et al., 2012; Jardine et al., 2013; McGuire et al., 2013). Following the priming immunization, Env immunogens designed to select for key antibody mutations can be administered in a specific order to guide antibody affinity maturation towards bnAb breadth and potency (Bonsignori et al., 2017; Bonsignori et al., 2016; Havenar-Daughton et al., 2018; Haynes et al., 2019; Haynes et al., 2012; Haynes et al., 2016; Huang et al., 2020; Saunders et al., 2019; Steichen et al., 2019; Zhang et al., 2016). However, guiding bnAb development is difficult because HIV-1 bnAbs are enriched in improbable functional somatic mutations that are required for neutralization potency and breadth (Bonsignori et al., 2017; Wiehe et al., 2018). Rare somatic mutations are due to the number of nucleotide changes needed for the amino acid substitution or the lack of targeting by the somatic mutation enzyme activation-induced cytidine deaminase (AID). To promote bnAb development, Envs will need to engage those B cell receptors that have accumulated functional improbable mutations, thereby selecting intermediate bnAb B cell lineage members to proliferate and evolve further (Bonsignori et al., 2017; Haynes et al., 2012; Wiehe et al., 2018). Whereas bnAbs arise in ∼50% of HIV-1 infected individuals (Hraber et al., 2014), to date, potent and durable bnAbs have not been induced in humans by vaccination. Together, these traits and roadblocks conspire to impede the easy induction of HIV-1 bnAbs.

HIV-1 Env is metastable and can adopt open and closed conformations (Tran et al., 2012; Ward and Wilson, 2017). Also, Env can be triggered by its cellular receptor, CD4, to open. The Env open conformation exposes non-neutralizing antibody (nnAb) epitopes that can create competition for Env antigen between nnAb and bnAb precursors (Havenar-Daughton et al., 2017; Lee et al., 2021; McGuire et al., 2014). To address the problem of Env trimers opening and the exposure of non-neutralizing epitopes, multiple strategies have been designed to stabilize Env trimers in native-like conformations (de Taeye et al., 2015; Guenaga et al., 2015; Henderson et al., 2020; Kong et al., 2016). We hypothesized that the inclusion of optimal stabilizing mutations will be critical for modified mRNAs to express antigenic and immunogenic Envs, since delivering immunogens directly as mRNA-LNP does not allow for immunogen purification. However, whether stabilizing mutations for modified mRNA expression of complex multimers will result in desired antigenicity and immunogenicity of trimer multimer NPs is not known.

We have previously demonstrated that a protein Env trimer designed with glycosylation sites eliminated in the first variable region (V1) of an autologous Env from an HIV-1 infected subject, CH848 (CH848 N133D N138T, CH848 10.17DT), conjugated to a ferritin nanoparticle was capable of initiating a V3-glycan bnAb lineage and selecting for key improbable mutations in immunized bnAb UCA heavy and light chain variable regions (VH + VL) knock-in (KI) mice (Saunders et al., 2019).

Here, we determined stabilization mutations in the CH848 10.17DT immunogen for formulation as mRNA-LNP. We demonstrate the modes of Env stabilization such that modified mRNA Env expression results in preferential binding to bnAbs of stabilized HIV-1 Envs in the forms of transmembrane gp160s, soluble gp140 SOSIP trimers, or gp140 SOSIP trimers on the surface of ferritin NPs encoded as a single-chain fusion gene mRNA (trimer-ferritin NPs). Moreover, we demonstrate that immunization of bnAb UCA VH + VL KI mice with mRNA-LNP encoding CH848 10.17DT gp160s or trimer-ferritin NPs initiate a V3-glycan bnAb B cell lineage, select for bnAb lineage B cells with BCRs bearing functional improbable mutations and induce high serum titers of tier 2 V3-glycan bnAb N332-dependent autologous neutralizing antibodies. Monoclonal antibodies (mAbs) from CH848 10.17DT trimer-ferritin NP mRNA-LNP vaccinated bnAb UCA VH+ VL KI mice acquired functional bnAb lineage improbable mutations and neutralized heterologous HIV-1 isolates. Thus, the modified mRNA-LNP vaccine platform can be used to encode complex scaffolded HIV-1 trimer multimer immunogens and initiate HIV-1 bnAb maturation.

## RESULTS

### Stabilization strategies for modified mRNA-encoded CH848 10.17DT Envs

Strategies have been proposed either to stabilize the Env trimer protein in the prefusion closed conformation or to prevent CD4-triggered structural rearrangements (de Taeye et al., 2015;Guenaga et al., 2015; Henderson et al., 2020; Kong et al., 2016; Zhang et al., 2018). We studied nine stabilization designs in CH848 10.17DT Env for expression as modified mRNAs (**Table S1**). The amino acid positions of these mutations are mapped onto the structure of CH848 10.17DT Env SOSIP trimer in **Figure 1A**.

**Figure 1.**
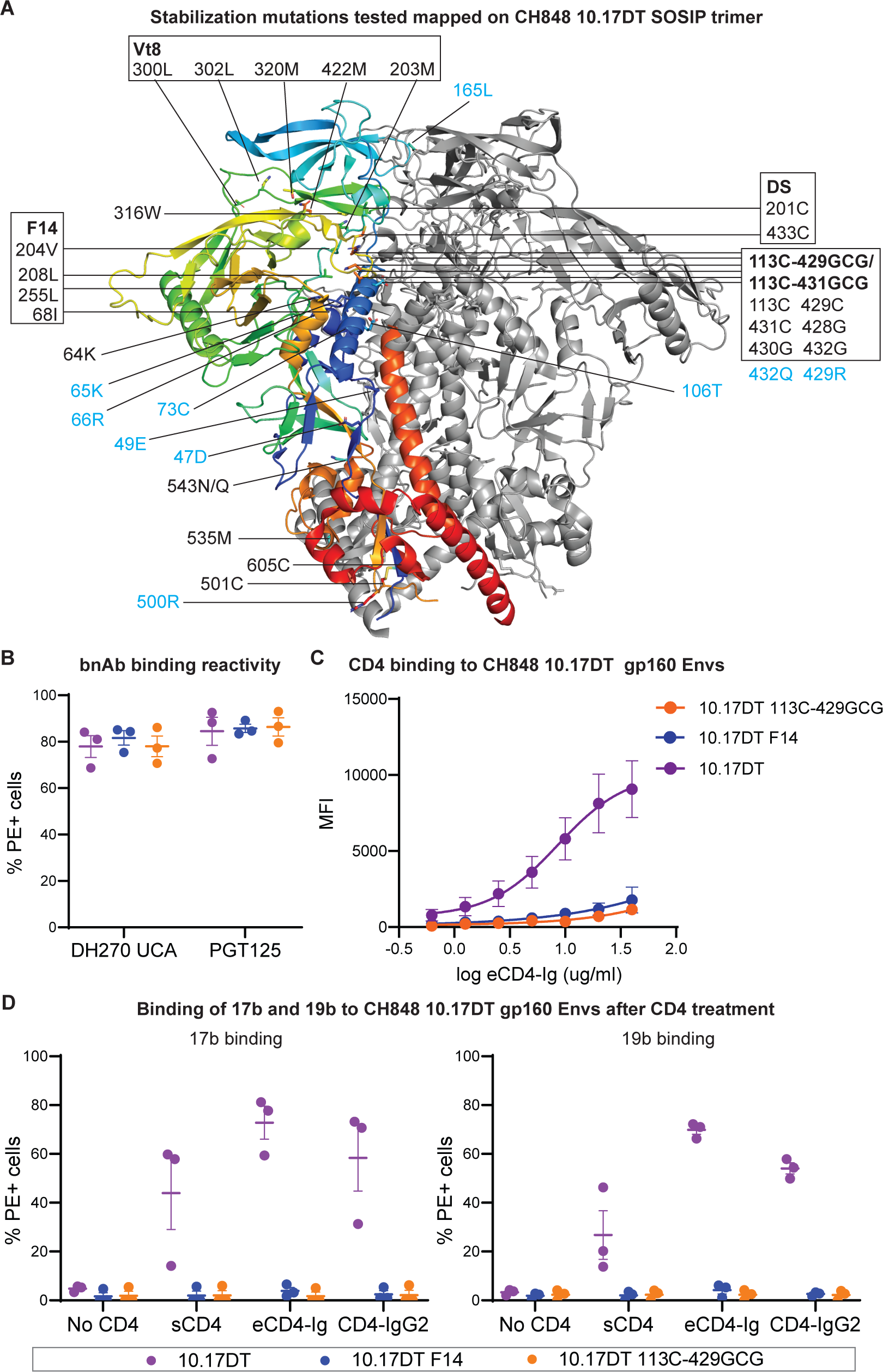
Antigenicity of modified mRNA-encoded CH848 10.17DT gp160 with stabilizing mutations. **(A)** Amino acid positions of stabilizing mutations tested in this study mapped onto structure of CH848 10.17DT SOSIP trimer (PDB ID: 6UM5). One protomer is shown in rainbow color and the other two protomers are shown in grey. Amino acid mutations in each stabilizing strategy are listed in boxes. Black fonts outside of boxes are mutations in v4.1, and blue fonts are mutations in v5.2.8, in addition to v4.1 mutations. Residue 561C in v5.2.8 or redesigned HR1 region in UFO mutation is not shown due to lack of HR1 region in this structure. **(B)** Antigenicity of modified mRNA-expressed CH848 10.17DT (purple), CH848 10.17DT F14 (blue), and CH848 10.17DT 113C-429GCG (orange) transmembrane gp160s measured by binding of V3-glycan bnAb PGT125 and the unmutated common ancestor of DH270 bnAb (DH270 UCA). Data were shown as means ± standard error of mean (SEM) of PE+ cell percentage among live cell population from three independent experiments. **(C)** eCD4-Ig binding reactivity to modified mRNA-expressed CH848 10.17DT, CH848 10.17DT F14, and CH848 10.17DT 113C-429GCG gp160s measured by flow cytometry. Data shown are means of mean fluorescent intensity (MFI) from three independent experiments. Error bars, mean ± SEM. **(D)** Binding reactivity of non-neutralizing antibodies (nnAbs) 17b (left) and 19b (right) to modified mRNA-expressed CH848 10.17DT, CH848 10.17DT F14, and CH848 10.17DT 113C-429GCG gp160s with or without treatment with sCD4, eCD4-Ig, and CD4-IgG2. Binding of nnAbs was shown as means ± SEM of PE+ cell percentage among live cell population from three independent experiments. See also Figure S1 and Table S1.

The DS mutations (201C-433C) introduce a disulfide bond in the closed Env trimer and prevents CD4-triggered exposure of the CCR5 co-receptor binding site and the V3 loop (Kwon et al., 2015). The F14 mutations (68I, 204V, 208L, 255L) are designed based on a structure of BG505 SOSIP trimer complexed with BMS-626529, a small molecule that blocks soluble CD4 (sCD4)-induced Env rearrangements (Pancera et al., 2017) and stabilize the SOSIP trimer by decoupling the allosteric conformational changes triggered by CD4 binding (Henderson et al., 2020). Vt8 mutations (203M, 300L, 302L, 320M, 422M) stabilize the V3 loop in the prefusion, V1/V2-coupled state (Henderson et al., 2020). The 113C-429GCG (113C-429C, 428G, 430G) and 113C-431GCG (113C-431C, 430G, 432G) mutations link the Env gp120 subunit inner and outer domains through a neo-disulfide bond, resulting in prefusion stabilized Env trimer with impaired CD4 binding (Zhang et al., 2018). For soluble gp140 trimer stabilization, we also tested SOSIPv4.1, v5.2.8 and uncleaved prefusion-optimized (UFO) mutations. Mutations in v4.1 (501C- 605C, 559P, R6, ΔMPER, 535M, 543N/Q, 316W, 64K) introduce hydrophobic amino acids to disfavor solvent exposure of the V3 loop and modify gp41 in the SOSIP.664 trimer, which improve trimer formation and thermostability and decrease V3 loop exposure (de Taeye et al., 2015). The UFO design replaces the bend between alpha helices in HR1 with a computationally designed linker and aims to minimize the metastability of HIV-1 gp140 trimer (Kong et al., 2016). Mutations in the v5.2.8 design (v4.1, 66R, 73C-561C, 165L, 432Q, 429R, 65K, 106T, 49E, 47D, 500R) are designed based upon v4.1 and combine an additional disulfide bond and eight trimer-derived mutations that stabilize BG505 SOSIP trimers (Guenaga et al., 2015). Both v4.1 and v5.2.8 include an improved hexa-arginine furin cleavage site R6 (Binley et al., 2002).

### Antigenicity of modified mRNA-encoded CH848 10.17DT gp160s with stabilizing mutations

We first designed modified mRNAs with stabilizing mutations encoding CH848 10.17DT Envs as transmembrane gp160s (**Table S1**) and tested their expression and antigenicity by transient transfection in Freestyle 293-F cells. All modified mRNA constructs expressed well and showed robust V3-glycan bnAb binding (**Figures 1B and S1**). In particular, CH848 10.17DT F14, CH848 10.17DT 113C-429GCG, and CH848 10.17DT 113C-431GCG gp160s exhibited binding reactivity to mature V3-glycan bnAb PGT125 and the DH270 UCA equal to that of CH848 10.17DT gp160 without stabilizing mutations (**Figures 1B and S1B**). The DS, Vt8, and F14/Vt8 mutations decreased DH270 UCA binding to CH848 10.17DT gp160s (**Figure S1B**). All CH848 10.17DT gp160s showed low binding to V2-glycan bnAbs PG9 and CH01 due to lack of a lysine at position 169 (K169) in the CH848 Env (McLellan et al., 2011). Thus, modified mRNA-encoded CH848 10.17DT F14, CH848 10.17DT 113C-429GCG, and CH848 10.17DT 113C-431GCG gp160s showed V3-glycan UCA antibody binding that is necessary for CH848 10.17DT germline targeting. To evaluate potential expression of CD4 induced (CD4i) non-neutralizing Env epitopes, we examined the susceptibility of each Env gp160 to CD4 triggering in transfected 293-F cells. Engineered (e) CD4-Ig (Fellinger et al., 2019) bound to CH848 10.17DT gp160 lacking stabilizing mutations in a dose-dependent manner, but binding of eCD4-Ig to CH848 10.17DT F14 and CH848 10.17DT 113C-429GCG gp160s was minimal (**Figure 1C**). Next, we assessed whether F14 or 113C-429GCG mutations could stabilize CH848 10.17DT gp160s in prefusion conformations and prevent the V3 loop or CCR5 co-receptor binding site exposure. Modified mRNA-transfected 293-F cells were either untreated or treated with 20 μg/ml of sCD4, eCD4-Ig (Fellinger et al., 2019) or CD4-IgG2 (Allaway et al., 1995), and Env conformation was determined by binding of CCR5 co-receptor binding site nnAb, 17b or distal V3 loop nnAb, 19b. In the absence of CD4 treatment, Env gp160s lacked binding to mAbs 17b and 19b (**Figures 1D and S1C**). After treatment with sCD4, eCD4-Ig, or CD4-IgG2, CH848 10.17DT gp160 without stabilizing mutations exhibited increased binding to both nnAbs 17b and 19b (**Figures 1D and S1C**). In contrast, stabilizing the Env gp160s with F14 or 113C-429GCG mutations completely prevented CD4- induced exposure of 17b and 19b epitopes (**Figures 1D and S1C**). Additionally, anti-gp41 nnAb 7B2 against the immunodominant epitope of gp41 (Pincus et al., 2003) showed low binding to modified mRNA-expressed CH848 10.17DT Env gp160s, confirming low exposure of this gp41 epitope (**Figure S1C**). Thus, CH848 10.17DT gp160s with F14 and 113C-429GCG mutations were stabilized such that they preferentially bound to bnAbs versus nnAbs and non-neutralizing epitope exposure after CD4 triggering was minimal.

### CH848 10.17DT gp160 mRNA-LNP elicited autologous tier 2 neutralizing antibodies in vivo

Based on stability and desired antigenicity, we selected CH848 10.17DT F14 and CH848 10.17DT 113C-429GCG gp160s to test their immunogenicity in heterozygous V3-glycan bnAb DH270 UCA heavy and light chains (VH^+/-^, VL+/- ) knock-in (DH270 UCA KI) mice (Saunders et al., 2019). Modified mRNAs encoding CH848 10.17DT F14 and CH848 10.17DT 113C-429GCG gp160s were encapsulated in ionizable LNP for immunization (**Figure 2A**). All mice immunized with CH848 10.17DT F14 or CH848 10.17DT 113C-429GCG gp160 mRNA-LNP developed serum binding IgGs to CH848 10.17DT trimer and gp120 monomer, and 3 CH848 10.17DT F14- and 4 113C-429GCG gp160 mRNA-LNP-vaccinated mice had IgGs binding to CH848 V3 peptide (**Figures 2B, S2A, and S2B**). Serum binding IgG titers to CH848 10.17DT trimer, gp120 monomer, or V3 peptide were not significantly different between CH848 10.17DT F14- and CH848 10.17DT 113C-429GCG-vaccinated groups one week after the third immunization (week 5) (**Figure 2B**, p > 0.05, Exact Wilcoxon Mann-Whitney U test), and serum binding to V3 peptide suggested exposure of the V3 loop *in vivo*. Importantly, we observed low to non-detectable levels of binding to gp41 in both groups of mice suggesting the gp41 was not exposed upon expression in vivo (**Figure S2D**). Both groups of mRNA-LNP immunizations induced mouse serum binding antibodies to CH848 10.17DT, CH848 10.17DT F14 and CH848 10.17DT 113C-429GCG gp160s on the surface of transfected 293-F cells (**Figure S2E**).

**Figure 2.**
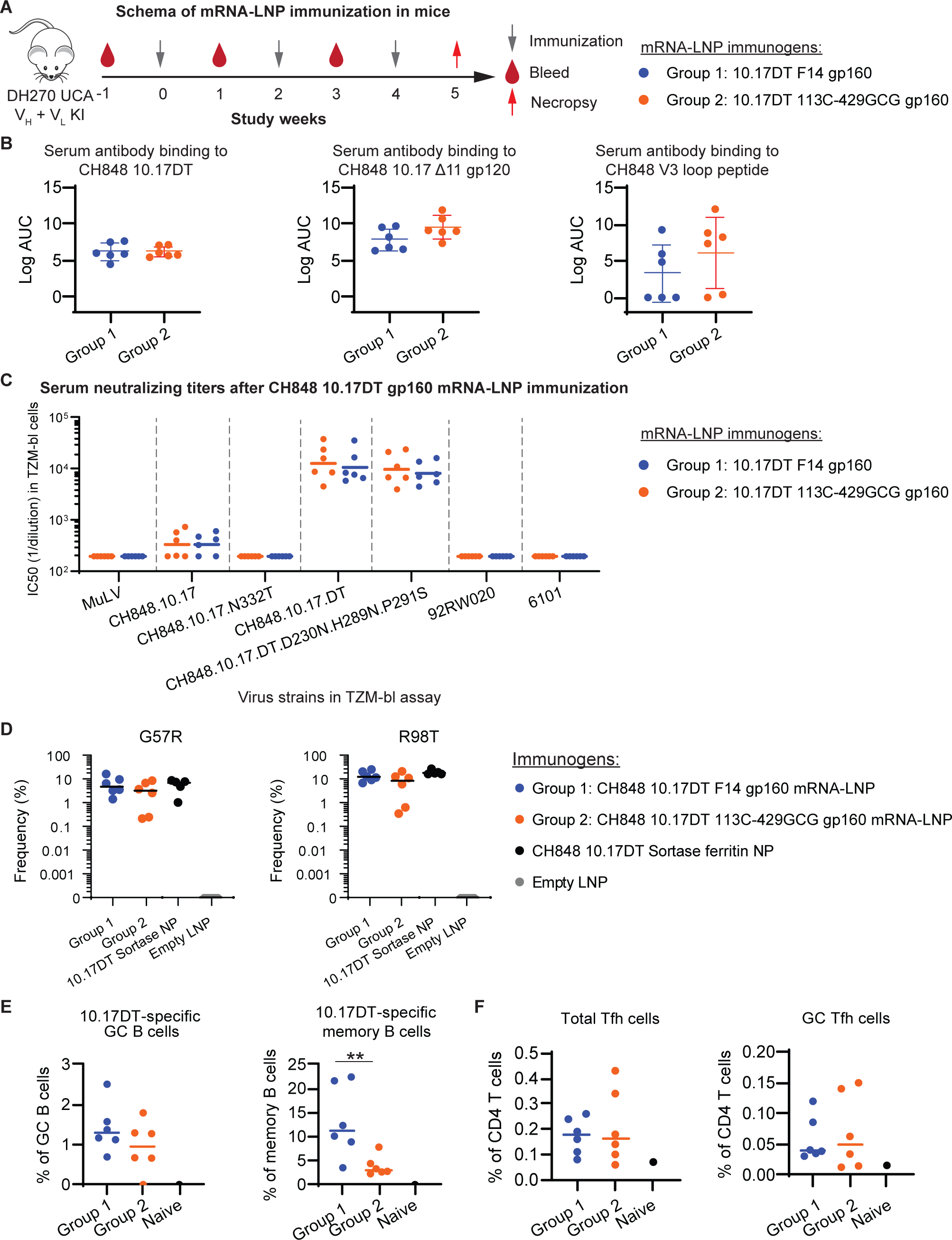
Immunogenicity of CH848 10.17DT gp160 mRNA-LNP in mice. **(A)** Immunization schema in DH270 UCA dKI mice with CH848 10.17DT F14 gp160 mRNA-LNP (blue) and CH848 10.17DT 113C-429GCG gp160 mRNA-LNP (orange). **(B)** Week 5 serum antibody binding to CH848 10.17DT SOSIP trimer, CH848 10.17 Δ11 gp120, and CH848 V3 loop peptide measured by ELISA. Data shown are log transformed area-under- curve (logAUC). Each dot represents an individual mouse (N = 6 each group). No significant statistical difference was observed between two groups (P > 0.05). **(C)** Week 5 serum neutralizing antibody titers measured in TZM-bl reporting cells with a panel of autologous and heterologous tier 2 HIV-1 pseudoviruses. Murine leukemia virus (MuLV) was used as negative control. Neutralization titers are reported as the serum dilution that inhibit 50% of virus replication (ID50). Each dot signifies an individual mouse (N = 6 each group). Horizontal bar indicates geometric mean titer (GMT) of ID50 in each group. No significant statistical difference was observed between two groups (P > 0.05). **(D)** Improbable G57R and R98T mutation frequencies in DH270 VH KI gene after CH848 10.17DT gp160 mRNA-LNP immunizations. Frequencies were compared to three CH848 10.17DT Sortase ferritin NP protein immunizations and empty LNP immunizations. Each dot represents an individual mouse (N = 6 in each group). Horizontal bar: median. **(E)** CH848 10.17DT gp160 mRNA-LNP vaccination induced GC B cell responses in spleen. Necropsy was performed on week 5 and splenocytes were subjected to GC responses immunophenotying by flow cytometry. Frequencies of CH848 10.17DT SOSIP trimer-specific GC B cells among total GC B cells (left) and CH848 10.17DT SOSIP trimer-specific memory B cells among total memory B cells (right) in splenocytes of CH848 10.17DT F14 gp160 mRNA-LNP (blue) or CH848 10.17DT 113C-429GCG gp160 mRNA-LNP (orange) vaccinated DH270 UCA dKI mice. Each dot represents an individual mouse (N = 6 in each group). Horizontal bar indicates means in each group. ** P < 0.01. **(F)** CH848 10.17DT gp160 mRNA-LNP vaccination induced GC Tfh cell responses in spleen. Frequencies of total Tfh (left) and GC Tfh cells (right) among CD4+ T cells in splenocytes at week 5 after CH848 10.17DT F14 gp160 mRNA-LNP or CH848 10.17DT 113C-429GCG gp160 mRNA- LNP vaccination in DH270 UCA dKI mice were assessed by flow cytometry. Each dot represents an individual mouse (N = 6 in each group). Horizontal bar indicates means in each group. No significant statistical difference was observed between two groups (P > 0.05). Significance was determined using Exact Wilcoxon Mann-Whitney U test. See also Figures S2 and S3.

Next, we asked whether CH848 10.17DT F14 and CH848 10.17DT 113C-429GCG gp160 mRNA-LNP immunizations in DH270 UCA KI mice elicited serum neutralizing antibodies. Neutralizing antibody titers one week after the third immunization (week 5) were assessed by the titration of sera needed to inhibit pseudovirus replication by 50% (ID50) in TMZ-bl reporter cells with a panel of 7 pseudotyped HIV-1 strains. As shown in **Figures 2C and S2F**, CH848 10.17DT F14 and CH848 10.17DT 113C-429GCG gp160 mRNA-LNP elicited autologous tier 2 (difficult-to-neutralize) (Mascola et al., 2005) neutralizing antibodies against CH848 10.17DT pseudovirus, with geometric mean titers (GMT) of ID50 at 13,175 and 10,820, respectively. Lower titers of neutralizing antibodies against CH848 10.17 virus with the V1 glycans restored (CH848 10.17) were induced that were N332 dependent, demonstrating targeting of the bnAb Env V3-glycan binding site. Moreover, comparable neutralization titers were observed against the CH848 10.17DT with mutations D230N H289N P291S designed to add glycans to occlude strain-specific, immunogenic regions on the Env, indicating that most neutralizing antibodies elicited by vaccination were not targeted to these glycan-bare regions (**Figures 2C and S2F**).

### CH848 10.17DT gp160 mRNA-LNP selected for key DH270 bnAb mutations and elicited germinal center responses

HIV-1 bnAbs are enriched in improbable functional somatic mutations in “cold-spots” of AID enzyme activity (Bonsignori et al., 2017; Wiehe et al., 2018). Splenocytes from one week after the third immunization (week 5) were subjected to next-generation sequencing (NGS) analysis. Both CH848 10.17DT F14 and CH848 10.17DT 113C-429GCG gp160 mRNA-LNP selected the critical improbable G57R mutation in the DH270 UCA VH KI gene that is necessary for the V3- glycan bnAb B cell lineage to acquire heterologous neutralization breadth (Bonsignori et al., 2017; Wiehe et al., 2018), with the medians of mutation frequency at 5.4% and 3.4%, respectively (**Figure 2D**). The antibodies also acquired a second key improbable VH R98T mutation (**Figure 2D**). Frequencies of the improbable VH G57R and the R98T mutations were comparable to those in a group of DH270 UCA KI mice immunized with Sortase ligated CH848 10.17DT Env ferritin NP protein (**Figure 2D**, P > 0.05). Thus, in DH270 UCA KI mice, CH848 10.17DT gp160 mRNA-LNP were immunogenic, induced potent N332-dependent autologous tier 2 neutralizing antibodies, and selected DH270 antibodies that acquired improbable mutations required for acquisition of heterologous HIV-1 neutralization (Bonsignori et al., 2017; Saunders et al., 2019; Wiehe et al., 2018).

To examine GC responses after CH848 10.17DT gp160 mRNA-LNP immunizations in DH270 UCA KI mice, splenocytes at week 5 were phenotyped for GC responses by flow cytometry using fluorophore-labeled CH848 10.17DT SOSIP trimer tetramers to detect CH848 10.17DT antigen-specific B cells (**Figure S3**). Both CH848 10.17DT F14 and CH848 10.17DT 113C-429GCG gp160 mRNA-LNP elicited CH848 10.17DT-specific GC B cells and memory B cells (**Figure 2E**). The average frequencies of CH848 10.17DT-specific GC B cells among total GC B cell population was 1.42% in CH848 10.17DT F14 gp160 mRNA-LNP group and 0.95% in CH848 10.17DT 113C-429GCG mRNA-LNP group. CH848 10.17DT F14 gp160 mRNA-LNP vaccinated group had higher frequencies of CH848 10.17DT-specific memory B cells among total memory B cells compared with CH848 10.17DT 113C-429GCG gp160 mRNA-LNP vaccinated group (mean at 13.14% versus 3.88%, P < 0.01, Exact Wilcoxon Mann-Whitney U test). Additionally, CH848 10.17DT gp160 mRNA-LNP elicited Tfh cell and GC Tfh cell responses in spleens in both groups (**Figures 2F and S3**).

### Antigenicity of modified mRNA-encoded CH848 10.17DT SOSIP trimers with stabilizing mutations

Next, antigenicity and stability of modified mRNA-encoded CH848 10.17DT SOSIP trimers with stabilizing mutations were evaluated (**Figure 1A and Table S1**). The CH848 10.17DT SOSIP trimers were chimeric with BG505 gp41 domain combined with the CH848 gp120 domain, upon which stabilizing mutations were added (Saunders et al., 2019). The antigenicity of modified mRNA-expressed *Galanthus nivalis* lectin (GNL)-purified CH848 10.17DT SOSIP trimers was measured by enzyme-linked immunosorbent assay (ELISA) using a panel of bnAbs and nnAbs. Each stabilized construct encoded by modified mRNA efficiently bound to the V3-glycan bnAbs 2G12, PGT125 and PGT128 and the DH270 lineage Abs DH270 UCA, DH270 IA4 and DH270.1 (**Figure 3A**). In particular, modified mRNA-encoded CH848 10.17DT SOSIP trimers with the DS mutations displayed greater binding reactivity to bnAbs, including DH270 lineage antibodies (DH270 UCA, DH270 IA4, and DH270.1) and cleaved trimer-specific gp41-gp120 interface bnAb PGT151, compared with other stabilizing mutations. Consistent with our observations with CH848 10.17DT gp160s, the Vt8 and F14/Vt8 mutations decreased DH270 UCA binding to CH848 10.17DT SOSIP trimers. CH848 10.17DT Vt8 and F14/Vt8 SOSIP trimers also displayed lower binding to trimer-specific bnAb PGT151 compared to CH848 10.17DT SOSIPv4.1, CH848 10.17DT DS, and CH848 10.17DT F14 SOSIP trimers, suggesting less native-like conformations of Envs with these latter mutations. Little to non-detectable binding to bnAbs was observed with v5.2.8 and UFO mutations combined (v5.2.8 + UFO).

**Figure 3.**
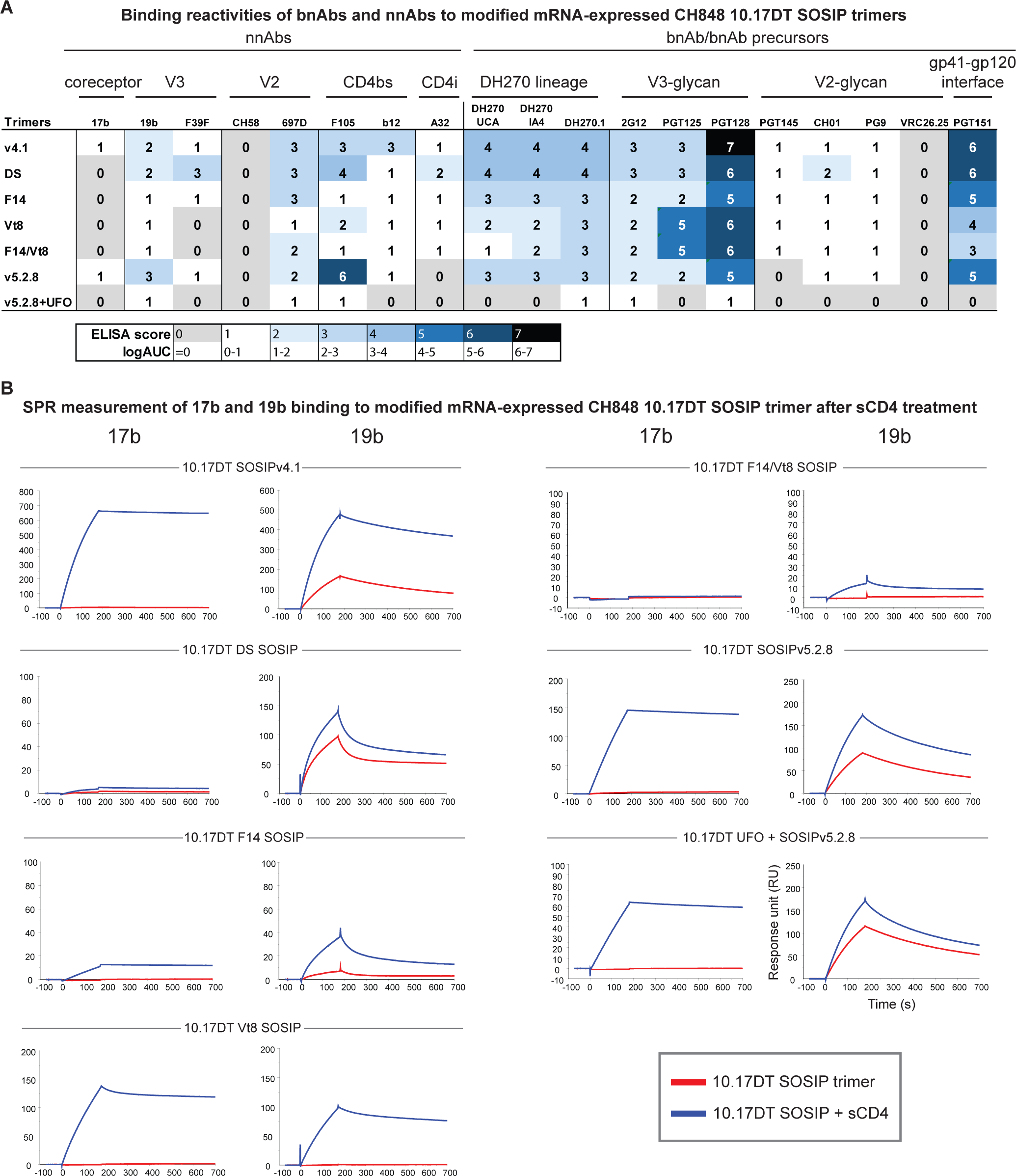
Antigenicity of modified mRNA-encoded CH848 10.17DT SOSIP trimers with stabilizing mutations. **(A)** BnAb/bnAb precursor and nnAb binding reactivity to modified mRNA-expressed CH848 10.17DT SOSIP trimers with various stabilizing mutations. Antibody binding was measured by ELISA. Data shown are means of logAUC from three independent experiments. **(B)** SPR sensorgrams of nnAb 17b or 19b binding to modified mRNA-expressed CH848 10.17DT SOSIP trimers with (blue) or without (red) sCD4 treatment. Antibodies 17b or 19b were immobilized onto a sensor chip. Modified mRNA-expressed GNL-purified CH848 10.17DT SOSIP trimers incubated with and without sCD4 were injected over the sensor chip surface. The protein was then allowed to dissociate for 600 seconds. See also Table S1.

All stabilized constructs tested, including CH848 10.17DT DS SOSIP trimers, presented low to non-detectable levels of binding to most nnAbs, except for CH848 10.17DT SOSIPv5.2.8 that displayed about 2-fold or higher binding to nnAbs 19b and F105, compared to other stabilized Envs tested (**Figure 3A**).

We assessed whether modified mRNA-expressed CH848 10.17DT SOSIP trimers with stabilizing mutations are resistant to CD4-induced opening by surface plasmon resonance (SPR). sCD4 treatment of modified mRNA-expressed non-stabilized CH848 10.17DT SOSIPv4.1 trimers increased binding of nnAb 17b (**Figure 3B**). In contrast, CH848 10.17DT DS, CH848 10.17DT F14, and CH848 10.17DT F14/Vt8 did not show binding to 17b with or without sCD4 treatment. Although CH848 10.17DT Vt8, CH848 10.17DT SOSIPv5.2.8, and CH848 10.17DT SOSIPv5.2.8+UFO trimers exhibited increased binding to 17b after sCD4 treatment, the binding was at a lower response level compared to CH848 10.17DT SOSIPv4.1. Similar trends were observed for 19b binding. An increase in 19b binding was observed with CH848 10.17DT SOSIPv4.1 trimer, while other constructs showed low levels of binding even after sCD4 triggering (**Figure 3B**). Thus, CH848 10.17DT DS when expressed by modified mRNA showed preferential binding to bnAbs with minimal exposure of non-neutralizing epitopes after CD4 treatment.

We next used size exclusion ultra-performance liquid chromatography (SE-UPLC) to define the folding of modified mRNA-encoded CH848 10.17DT SOSIP trimers. The analytical SE-UPLC profile of PGT151-purified CH848 10.17DT DS SOSIP trimer indicated that a well-folded CH848 10.17DT SOSIP trimer was separated and eluted from the column as shown in **Figure 4A**. GNL-purified modified mRNA-expressed CH848 10.17 DT SOSIPv4.1 and CH848 10.17DT DS SOSIP trimer samples showed a dominant peak of trimer that was 62% and 65% of the total peak, respectively (**Figures 4B and 4C**). As shown in **Figure 4D**, negative stain electron microscopy (NSEM) analysis of CH848 10.17DT DS trimer confirmed the expression of well-folded SOSIP trimers from modified mRNA-transfected 293-F supernatant. In summary, DS mutation was the optimal stabilizing mutation strategy for CH848 10.17DT Env SOSIP trimers.

**Figure 4.**
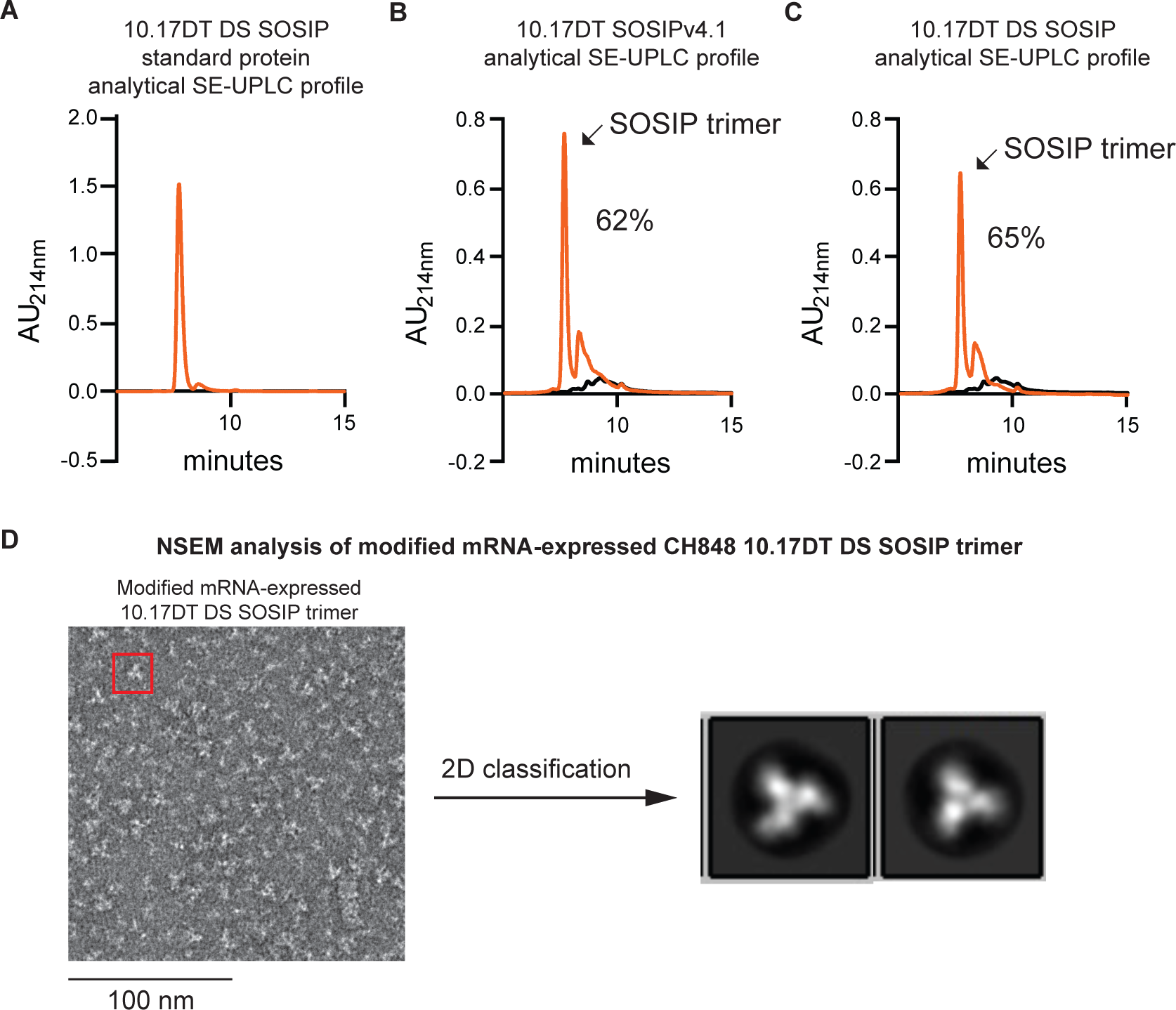
Modified mRNA-expressed CH848 10.17DT DS SOSIP trimer is well-folded. **(A)** Analytical size-exclusive ultra-performance liquid chromatography (SE-UPLC) profile of CH848 10.17DT SOSIP trimer protein standard purified by bnAb PGT151. CH848 10.17DT SOSIP trimer elutes from the column at about 7 min. **(B-C)** Analytical SE-UPLC profile of modified mRNA-expressed GNL-purified **(B)** CH848 10.17DT SOSIPv4.1 trimers (orange) and **(C)** CH848 10.17DT DS SOSIP trimers (orange). Black curve indicates GNL-purified material from mock transfection. **(D)** Negative-stain electron microscopy (NSEM) analysis of modified mRNA-expressed GNL-purified CH848 10.17DT DS SOSIP trimers. Shown on the left is a representative NSEM micrograph of CH848 10.17DT DS SOSIP trimer and on the right are 2D classification of well- folded trimers.

### Antigenicity of CH848 10.17DT SOSIP trimer-ferritin NPs with stabilizing mutations encoded by modified mRNAs

We recently demonstrated that sortase A-ligated CH848 10.17DT Env trimer ferritin NPs were potent priming immunogens for bnAb precursors (Saunders et al., 2019). Here we asked if SOSIP trimers with stabilizing mutations could be presented in an arrayed manner on ferritin and self-assemble into trimer-ferritin NPs when encoded by a single-chain modified mRNA. CH848 10.17DT SOSIP trimer-ferritin NPs were produced by gene fusion of CH848 10.17DT SOSIP trimer gene with *Helicobacter pylori* (*H. pylori)* ferritin gene (*FtnA*) (GenBank NP_223316) and were tested for expression, stability, and antigenicity (**Figure 5A**). We also constructed CH848 10.17DT SOSIP trimer-ferritin NPs with CH848 strain-specific, immunogenic regions occluded by adding glycans (CH848 10.17DT with D230N, H289N, P291S) in addition to adding the E169K mutation, which is critical to interactions with V2-glycan bnAbs, including trimer-specific bnAb PGT145 (Doria-Rose et al., 2012; Lee et al., 2017; McLellan et al., 2011). This CH848 10.17DT Env trimer (CH848 10.17DT D230N, H289N, P291S, E169K) was termed “enhanced CH848 10.17DT” (CH848 10.17DTe). Since the linker sequence connecting ferritin and Env protein would affect the expression and assembly of NPs, we tested CH848 10.17DTe DS trimer-ferritin NPs with two different linkers, the sequences of which were GGGSGGGGSGLSK (termed “2xGS linker”) and GGGSGGGGSGGGGSGLSK (termed “3xGS linker”). We also designed another trimer-ferritin fusion construct using the *H. pylori* ferritin with a N19Q mutation, which removed a potential N-linked glycosylation site at position 19 and added a glycine and a serine to the C-terminus of the ferritin protein (hereafter termed the “VRC ferritin”) (Kanekiyo et al., 2013).

**Figure 5.**
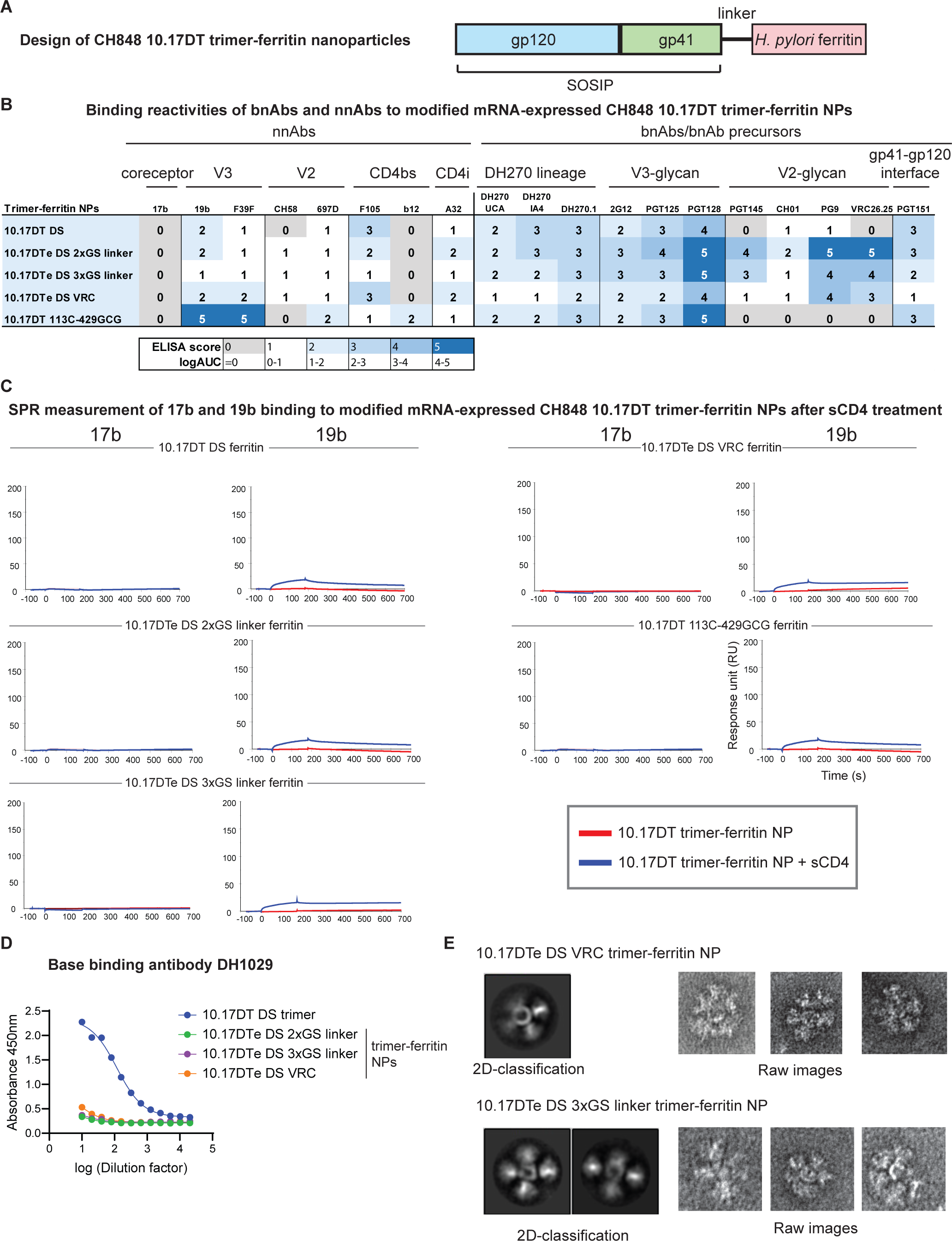
Antigenicity of CH848 10.17DT SOSIP trimer-ferritin NPs with stabilizing mutations. **(A)** Design of CH848 10.17DT SOSIP trimer-ferritin fusion NPs. The CH848 10.17DT SOSIP gene is genetically fused to *Helicobacter pylori* Ferritin gene (*FtnA*) by a linker sequence. Ferritin self- assembles into a 24-mer nanoparticle, with 8 SOSIP trimers on the surface. **(B)** BnAb/bnAb precursor and nnAb binding reactivity to modified mRNA-expressed CH848 10.17DT SOSIP trimer-ferritin NPs with various stabilizing mutations. Antibody binding was measured by ELISA. Data shown are means of logAUC from three independent experiments. **(C)** SPR sensorgrams of nnAb 17b or 19b binding to modified mRNA-expressed CH848 10.17DT SOSIP trimer-ferritin NPs with (blue) or without (red) sCD4 treatment. Antibodies 17b or 19b were immobilized onto a sensor chip. Modified mRNA-expressed GNL-purified CH848 10.17DT SOSIP trimer-ferritin NPs incubated with and without sCD4 were injected over the sensor chip surface. The protein was then allowed to dissociate for 600 seconds. **(D)** Binding reactivity of Env base antibody DH1029 to modified mRNA-expressed CH848 10.17DT DS SOSIP trimer (blue) or CH848 10.17DTe DS 2xGS linker (green), CH848 10.17DTe DS 3xGS linker (purple), and CH848 10.17DTe DS VRC (orange) trimer-ferritin NPs measured by ELISA. Data shown are means of absorbance at 450nm from at least two independent experiments. **(E)** Representative NSEM images (right) and 2D classifications (left) of modified mRNA- expressed, PGT145-purified CH848 10.17DTe DS VRC trimer-ferritin NPs (top) and CH848 10.17DTe DS 3xGS linker trimer-ferritin NPs (bottom). See also Figure S4 and Table S1.

All CH848 10.17DT and CH848 10.17DTe trimer-ferritin NPs exhibited effective binding to V3- glycan bnAbs tested (**Figure 5B**). CH848 10.17DT DS and CH848 10.17DT 113C-429GCG trimer-ferritin NPs without the E169K mutation, as expected, displayed weak or no binding to V2- glycan bnAbs PGT145, CH01, PG9, and VRC26.25. Thus, modified mRNAs with stabilizing strategies tested were able to encode 10.17DT SOSIP trimer-ferritin NPs that were antigenic for bnAbs when expressed *in vitro*. In contrast, all CH848 10.17DT and CH848 10.17DTe trimer-ferritin NPs showed low to non-detectable binding to nnAbs (**Figure 5B**). Specifically, none of the trimer-ferritin NPs showed binding to 17b, and the binding to 19b was low, except for CH848 10.17DT 113C-429GCG trimer-ferritin NP, indicative of an exposed distal V3 loop.

SPR analysis following sCD4 treatment showed no increased binding of nnAb 17b to CH848 10.17DT and CH848 10.17DTe SOSIP trimer-ferritin NPs and low levels of binding of nnAb 19b (**Figure 5C**). Thus, we demonstrated that CH848 10.17DT trimer-ferritin NPs could be expressed with modified mRNAs and bound to bnAbs efficiently with limited binding to nnAbs when optimized stabilizing mutations were present. Additionally, the base part of Env trimer proteins has been shown to be highly immunogenic and some base binding antibodies can disassemble Env trimers into monomers and cause the exposure of nnAb epitopes (Turner et al., 2021). Thus, we tested binding of CH848 10.17DT trimer-ferritin NPs to an Env base binding antibody DH1029. Modified mRNA-expressed CH848 10.17DT DS SOSIP trimers without ferritin bound to DH1029 strongly, while no binding was observed to modified mRNA-expressed CH848 10.17DT trimer-ferritin NPs (**Figure 5D**), demonstrating that the immunodominant base of Env trimers was not accessible to base binding antibody DH1029 recognition when presented as a multimeric nanoparticle on ferritin.

Next, we assessed if modified mRNA-expressed CH848 10.17DT trimer-ferritin protein indeed self-assembled into NPs by NSEM. We purified modified mRNA-transfected 293-F supernatants of CH848 10.17DT DS VRC and CH848 10.17DT DS 3xGS linker trimer-ferritin NPs by PGT145 and demonstrated that CH848 10.17 DTe DS VRC ferritin and 3xGS linker ferritin mRNA transfection produced stabilized Env trimer-ferritin NPs (**Figures 5E and S4**, yellow circles). Few free trimers were observed (**Figure S4**, purple arrow). Host protein particles were classified into small 7-fold symmetry particles (**Figure S4**, yellow arrow) and large polygon-shaped particles (**Figure S4**, red arrow). The 7-fold symmetry particles were compatible with proteasomes (Adams, 2003). Polygon-shaped particles have been observed in HIV-1 Env protein preparation by others (He et al., 2016), and are compatible with secreted Galectin-3 binding proteins (Gal-3BP) that assemble into ring-like polymers (Muller et al., 1999; Sasaki et al., 1998), and were co-purified with HIV-1 Env NPs. Thus, NSEM analysis demonstrated that CH848 10.17DT trimer-ferritin fusion proteins self-assembled into well-folded NPs.

### CH848 10.17DT SOSIP trimer-ferritin NP mRNA-LNP induced autologous tier 2 neutralizing antibodies

To assess the immunogenicity of mRNA-LNP encoding CH848 10.17 DT trimer-ferritin NPs, we immunized DH270 UCA KI mice (**Figure 6A**). All CH848 10.17DT trimer-ferritin NP mRNA-LNP elicited serum antibody bound to CH848 10.17DT and CH848 Δ11 gp120 proteins (**Figures 6B, S5A, and S5B**). Interestingly, in contrast to CH848 10.17DT gp160s (**Figure 2B**), none of the trimer-ferritin NPs induced V3 loop peptide binding antibodies, suggesting that trimer-ferritin NPs had greater stabilization of the V3 loop (**Figures 6B and S5C**). To address the concern that using *H. pylori* ferritin induces antibodies that target the ferritin protein itself, we tested serum antibody binding to *H. pylori* ferritin used in our NPs and to human ferritin protein. We detected binding activity to *H. pylori* ferritin but did not observe any immunized mouse serum cross-reactivity with human ferritin (**Figures S6E and S6F**). To assess whether trimer base-binding antibodies were elicited, we determined if immunized mouse serum contained antibodies that could block the trimer base-binding antibody DH1029. No blocking of DH1029 binding was observed in CH848 10.17DT trimer-ferritin NP mRNA-LNP vaccinated mice, except for one mouse vaccinated with CH848 10.17DT 113C-429GCG trimer-ferritin NP mRNA-LNP (background cut-off at 20%). In contrast, sera from a control group of CH848 10.17DT DS SOSIP trimer protein vaccinated DH270 UCA KI mice showed DH1029 blocking activity after the second and third immunizations (**Figure 6C**). Thus, vaccination with CH848 10.17DT trimer-ferritin NP mRNA-LNP in DH270 UCA KI mice did not elicit trimer base-targeted antibodies whereas CH848 10.17DT DS SOSIP trimer protein did elicit trimer base off target antibodies.

**Figure 6.**
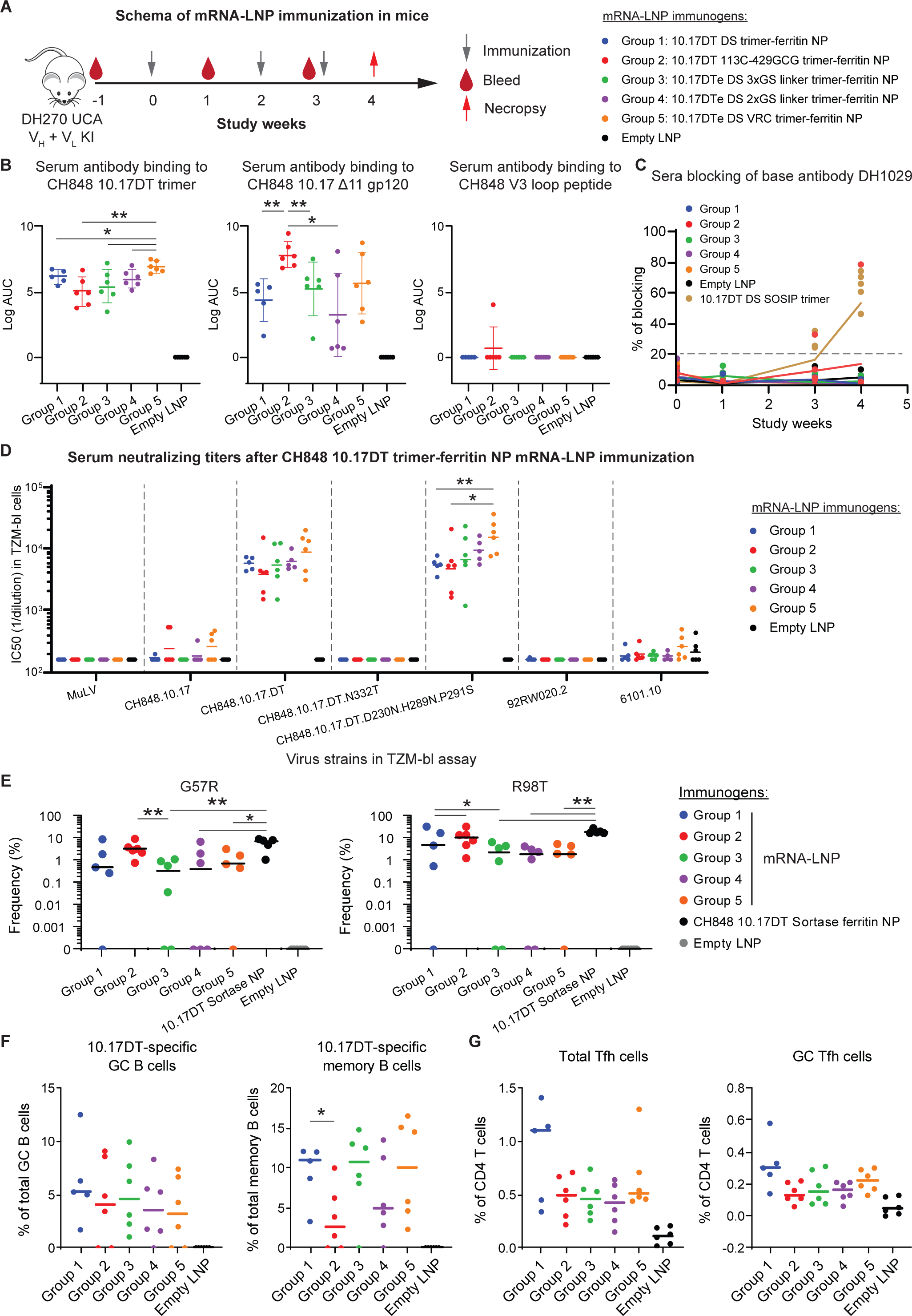
Immunogenicity of CH848 10.17DT SOSIP trimer-ferritin NP mRNA-LNP in mice. **(A)** Immunization schema with CH848 10.17DT SOSIP trimer-ferritin NP mRNA-LNP in DH270 UCA dKI mice. **(B)** Serum IgG binding to CH848 10.17DT SOSIP trimer, CH848 10.17 Δ11 gp120, and CH848 V3 peptide. Each dot represents an individual mouse (N = 5 in Group 1; N = 6 in the rest of groups). * P < 0.05, ** P < 0.01. **(C)** DH1029 blocking by sera from mice vaccinated with CH848 10.17DT trimer-ferritin NP mRNA- LNP. CH848 10.17DT trimer protein vaccinated mice from one of our previous studies were included as a positive control (brown). Dotted horizontal line indicates background cut-off at 20% blocking. **(D)** Serum neutralizing titers after CH848 10.17DT SOSIP trimer-ferritin NP mRNA-LNP immunizations measured in TZM-bl reporter cells. MuLV was used as negative control. Neutralization titers are reported as the serum dilution that inhibit 50% of virus replication (ID50). Each dot signifies an individual mouse (N = 5 in Group 1; N = 6 in the rest of groups). Horizontal bar indicates geometric mean titer (GMT) of ID50. * P < 0.05, ** P < 0.01. **(E)** Improbable G57R and R98T mutations frequency in DH270 VH KI gene after CH848 10.17DT SOSIP trimer-ferritin mRNA-LNP immunizations. Frequencies were compared to three CH848 10.17DT Sortase ferritin NP protein immunizations and empty LNP immunizations. * P < 0.05, ** P < 0.01. **(F)** Frequencies of CH848 10.17DT-specific GC B cells and memory B cells in splenocytes of CH848 10.17DT SOSIP trimer-ferritin mRNA-LNP vaccinated mice. * P < 0.05 **(G)** Frequency of CH848 10.17DT-specific GC Tfh cells and memory B cells in spleen of CH848 10.17DT SOSIP trimer-ferritin mRNA-LNP vaccinated mice. * P < 0.05 Significance was determined by Exact Wilcoxon Mann-Whitney U test, without any P value adjustment for multiple comparison. See also Figures S5 and S3.

Next, we assessed tier 2 serum neutralizing antibody titers after 3 immunizations against a panel of HIV-1 strains in the TZM-bl neutralization assay. All CH848 10.17DT SOSIP trimer-ferritin NP mRNA-LNP elicited neutralizing antibodies against autologous tier 2 virus CH848 10.17DT in an N332-dependent manner (**Figures 6D and S6G**). Comparable neutralizing titers against glycan holes-filled CH848 10.17DT virus (230N, 289N, 291S) were observed, indicating the antibody responses were not directed at glycan holes but rather were targeted to the V3-glycan bnAb site. CH848 10.17DTe DS VRC trimer-ferritin NP mRNA-LNP vaccinated mice showed higher neutralizing titers to CH848 10.17DT 230N, 289N, 291S viruses compared to CH848 10.17DT DS (P < 0.05, Exact Wilcoxon Mann-Whitney U test) and CH848 10.17DT 113C- 429GCG (P < 0.01, Exact Wilcoxon Mann-Whitney U test) trimer-ferritin NP mRNA-LNP vaccinated groups. The CH848 10.17DTe DS VRC trimer-ferritin NPs induced both the highest binding to CH848 10.17DT trimer and the highest level of tier 2 neutralizing antibodies to CH848 10.17 DT and the glycan holes filled virus version (CH848 10.17DT D230N, H289N, P291S) (**Figure 6D**). Thus, CH848 10.17DT SOSIP trimer-ferritin NPs encoded as mRNA-LNP efficiently elicited tier 2 autologous neutralizing antibodies that targeted the N332-dependent V3 glycan bnAb site.

### CH848 10.17DT SOSIP trimer-ferritin NP mRNA-LNP elicited germinal center responses and selected for key DH270 bnAb mutations

All five CH848 10.17DT trimer-ferritin NPs selected the improbable VH G57R mutation in DH270 UCA KI gene, with the highest median of mutation frequency at 3.2% observed in CH848 10.17DT 113C-429GCG trimer-ferritin NP vaccinated mice. Similarly, the R98T mutation in the DH270UCA VH KI gene was also selected in all groups by mRNA-LNP (**Figure 6E**). Thus, mRNA-LNP encoded CH848 10.17DT trimer-ferritin NP immunizations in DH270 UCA KI mice efficiently elicited key improbable and other mutations in DH270 intermediate antibodies.

All five CH848 10.17DT trimer-ferritin NP mRNA-LNP induced CH848 10.17DT-specific GC B cells and memory B cells in spleens (**Figure 6F**). Tfh cells and GC Tfh cells were also observed in all CH848 10.17DT SOSIP trimer-ferritin NP vaccinated mice (**Figure 6G**). No significant difference was observed among 5 immunization groups (P < 0.05, Exact Wilcoxon Mann-Whitney U test). Interestingly, empty LNP immunizations also elicited Tfh cell responses, albeit at much lower frequencies, consistent with previous observations that mRNA-LNP may have adjuvant effects that favors Tfh cell and GC responses (Pardi et al., 2018a).

### CH848 10.17DT trimer-ferritin NP mRNA-LNP immunization induced heterologous neutralizing mAbs that acquired improbable mutations

To further assess antibody responses elicited by CH848 10.17DT trimer-ferritin NP mRNA-LNP, we injected DH270 UCA KI mice intradermally (i.d.) or intramuscularly (i.m.) with CH848 10.17DT DS trimer-ferritin NP modified mRNA-LNP for six immunizations and sorted CH848 10.17DT-specific single memory B cells on 96-well plates and amplified immunoglobulin (Ig) heavy and light chain variable regions by PCR (**Figures 7A and S5A**). We cloned a total of 397 Ig heavy and light chain pairs, 228 (57%) pairs of which used DH270 KI genes *IGVH1-2* and *IGVL2-23*. Among these 228 DH270-like antibodies, 173 (76%) antibodies have acquired at least one amino acid mutation (**Figures 7B and S5B**). We aligned all unique VH1-2/VL2-23 Ig gene amino acid sequences with bnAb DH270.6, and found that the Ig heavy chain group accumulated a total of 14 out of 19 (74%) DH270.6 probable mutations and 4 out of 8 (50%) DH270.6 improbable mutations (**Figure S7**). Similarly, the Ig light chain group accumulated 6 out of 9 (67%) DH270.6 probable mutations and 4 out of 6 (67%) improbable mutations (**Figure S8**). Specifically, 5 (2%) and 20 (9%) Ig heavy chains acquired the DH270.6 bnAb G57R and the R98T improbable mutations, respectively; and 2 (1%) Ig light chains acquired the DH270.6 bnAb L48Y improbable mutation (**Figure 7C**). Binding reactivity of cloned antibodies were screened in ELISA (**Table S3**), and antibodies with heterologous HIV-1 A.Q23 Env binding were selected for further assessment. Among them, three mAbs (DH270.mo84, DH270.mo85, and DH270.mo86) were identified that showed strong binding to CH848 10.17DT, CH848 10.17, and heterologous HIV-1 A.Q23 SOSIP trimers (**Figure 7D**). We then assessed neutralization of these three mAbs against a panel of 17 HIV-1 isolates that the first intermediate ancestor antibody (DH270 IA4) in the DH270 lineage neutralizes (Saunders et al., 2019) (**Figure 7D**). Each of the 3 mAbs neutralized autologous tier CH848 viruses and heterologous 92RW020 and 6101.1 viruses with titers comparable to DH270 IA4 (Saunders et al., 2019). Antibody DH270.mo84 also neutralized each of the 13 other heterologous HIV-1 isolates tested. Antibodies DH270.mo85 and DH270.mo86 neutralized 8 and 10 of 13 heterologous isolates, respectively (**Figure 7D**). DH270.mo84 encoded the VH G57R improbable mutation, DH270.mo86 encoded the VH R98T improbable mutation, while DH270.mo85 had both the G57R and R98T improbable mutations. Additionally, DH270.mo86 acquired VL S27Y and VL S57N improbable mutations (**Figure 7E**). Thus, CH848 10.17DT DS trimer-ferritin NP mRNA-LNP immunization induced heterologous tier 2 neutralizing DH270 antibodies with improbable mutations.

**Figure 7.**
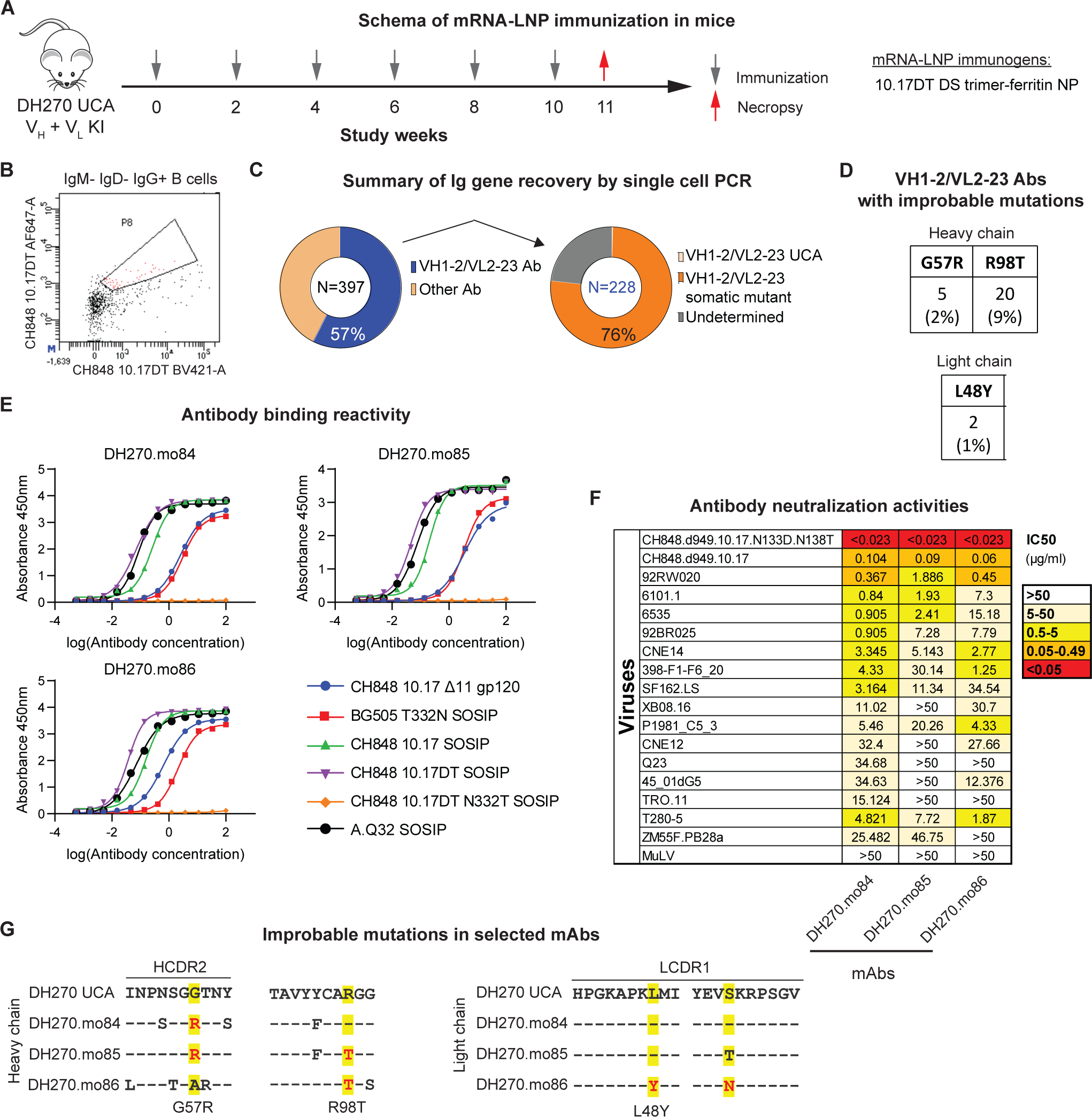
CH848 10.17DT DS trimer-ferritin NP mRNA-LNP vaccination elicited antibodies that acquired improbable mutations and neutralization breadth. **(A)** Immunization schema with CH848 10.17DT DS SOSIP trimer-ferritin NP mRNA-LNP in DH270 UCA dKI mice. Necropsy was performed one week after the sixth immunization. **(B)** Representative gate for CH848 10.17DT Env-specific single memory B cell sorting from DH270 UCA dKI mice splenocytes one week after the sixth vaccination with CH848 10.17DT DS trimer-ferritin NP mRNA-LNP. **(C)** Summary of Ig gene recovery by PCR from single-cell sorted memory B cells. Left: A total of 397 Ig heavy and light chain gene pairs were recovered. 228 (57%) of cloned Ig gene pairs used DH270 *IGVH1-2* and *IGVL2-23* genes, and thus were considered as DH270-like Abs. Ig gene pairs that used only one of DH270 heavy chain or light chain, or endogenous mouse Ig genes are categorized as “Other Ab”. Right: The number and percentage of the 228 DH270-like VH1-2/VL2- 23 Abs that had acquired at least one amino acid change in heavy or light chain (N = 173, 76%). VH1-2/VL2-23 Abs without full length, clean VDJ/VJ sequences are categorized as “Undetermined”. **(D)** Summary of improbable mutations in VH1-2/VL2-23 Abs. Among the 228 DH270-like VH1- 2/VL2-23 Abs, 5 (2%) and 20 (9%) of them have acquired the VH G57R and the R98T improbable mutations, respectively; 2 (1%) have acquired the VL L48Y improbable mutation. **(E)** Binding reactivity of three cloned mAbs DH270.mo84, DH270.mo85, and DH270.mo86 to a panel of CH848 10.17DT proteins measured by ELISA. All three antibodies bound to CH848 10.17DT SOSIP trimers and CH848 10.17 SOSIP trimers, while no binding to CH848 10.17DT N332T was observed. Binding to BG505 T332N and heterologous Q23 SOSIP trimers was also observed. **(F)** DH270.mo84, DH270.mo85, and DH270.mo86 neutralization activity against a panel of 17 HIV-1 isolates. Data shown are antibody concentration that inhibit 50% of virus replication (IC50) in TZM-bl assay. MuLV was used as negative control. **(G)** Improbable mutations in mAbs DH270.mo84, DH270.mo85, and DH270.mo86. Alignment of mAb heavy and light chain sequences with DH270 UCA sequences. Mutations are highlighted and improbable mutations are shown in red fonts. See also Figures S5-S7 and Table S3.

## DISCUSSION

Nucleoside-modified mRNAs in lipid nanoparticles (mRNA-LNP) represent an exciting new platform for viral vaccine development for experimental HIV-1 vaccines (Baden et al., 2020; Mu et al., 2021; Pardi et al., 2018b; Polack et al., 2020). A major question is if mRNA designs should incorporate stabilizing mutations in trimers or NPs to optimize immunogen expression and stability since the protein products of mRNAs cannot be purified after mRNA-LNP injection *in vivo*. In this study, we evaluated mutations that stabilize modified mRNA-encoded Envs expressed as transmembrane gp160s, soluble SOSIP trimers, or single-gene mRNA trimer-ferritin NPs. For mRNAs encoding the V3-glycan germline targeting CH848 10.17DT Env (Saunders et al., 2019), we showed that F14 and 113C-429GCG mutations optimally stabilized transmembrane gp160s, DS mutation best stabilized SOSIP trimers, and were also able to optimally stabilize trimer-ferritin NPs expressed from modified mRNA.

Since we have previously demonstrated that immunization with Env ferritin NPs is superior to soluble trimers alone (Saunders et al., 2019), we determined immunogenicity of mRNA-LNP encoding transmembrane Env gp160 and trimer-ferritin NPs for their ability to expand UCAs of V3-glycan DH270 bnAb B cell lineage and to select for desired, functional bnAb mutations. mRNA-LNP encoding both transmembrane gp160s and soluble trimer-ferritin NPs induced high titers of autologous bnAb-targeted tier 2 neutralizing antibodies with groups of mutations present in bnAb intermediate antibodies (Bonsignori et al., 2017; Saunders et al., 2019). Additionally, mAbs with heterologous neutralizing activities and functional improbable mutations were isolated after trimer-ferritin NP mRNA-LNP vaccination. Moreover, we observed accumulation of mature bnAb DH270.6 mutations in vaccine-induced mAbs, although mutations are distributed across all isolated mAbs. This observation suggests that mRNA-LNP immunogens are selecting for key bnAb mutations, so the next goal is to have mutations concatenated on one or two mAbs. To achieve this, prolonged GC responses or enhanced recruitment of memory B cells back into GCs will likely need to be induced by vaccination to allow bnAb lineage B cell BCRs to acquire more mutations.

Eliciting bnAbs by vaccination has not been successful. However, studies in HIV-1-infected individuals have demonstrated that those who make bnAbs have higher levels of T follicular helper (Tfh) cells (Locci et al., 2013; Moody et al., 2016), NK cell dysfunction (Bradley et al., 2018), defects in T regulatory cells (Treg) (Moody et al., 2016), and B cell repertoires containing longer HCDR3-bearing B cells and autoreactive B cells that normally are deleted (Roskin et al., 2020).

Nucleoside-modified mRNA-LNP vaccines selectively induce high levels of Tfh cells and minimize induction of Treg cells (Pardi et al., 2018a), and thus will be a key platform for bnAb lineage initiation and selection of B cells with improbable functional mutations that facilitate bnAb maturation. The CH848 10.17DTe DS trimer-ferritin NP is currently in good manufacturing practice (GMP) production to investigate the priming of such lineages in humans both as mRNA-LNP or recombinant protein.

In summary, we have demonstrated that single-chain mRNAs can be designed to encode complex molecules such as HIV-1 Env trimer-ferritin NP and that these immunogens are capable of selecting for difficult-to-elicit improbable mutations critical for broad tier 2 virus neutralization. The complex biology of HIV-1 bnAbs necessitates a vaccine strategy that utilizes a series of sequentially administered Env immunogens that initially expand bnAb precursors and then select for improbable mutations (Haynes et al., 2019; Haynes et al., 2012; Saunders et al., 2019; Wiehe et al., 2018). Manufacturing of complex nanoparticle protein immunogens in large-scale is faced with significant practical and funding challenges. The use of mRNA-LNP has the possibility of making such a complex immunization regimen both logistically achievable and potentially cost- effective.

## ACKNOWLEGEMENTS

We thank Holly Zoeller for assistance with SE-UPLC assays. We thank Cindy Bowman, Grace Stevens, and Austin Harner for help with animal studies. We thank Victoria Gee-Lai and Maggie Barr for help with ELISA assays. We also thank Cynthia Nagel for project management. Flow cytometry was performed in the Duke Human Vaccine Institute Research Flow Cytometry Facility (Durham, NC). Surface Plasmon Resonance was performed in the Biacore core facility at Duke Human Vaccine Institute. Next-generation sequencing was performed at the Duke Human Vaccine Institute Viral Genetic Analysis core facility. This project was supported by NIH, NIAID, Division of AIDS Intergrated Preclinical and Clinical AIDS Vaccine Development Grant AI135902, and by NIAID, Division of AIDS Consortia for HIV/AIDS Vaccine Development (CHAVD) Grant UM1AI144371.

## AUTHOR CONTRIBUTIONS

Conceptualization, Z.M., K.O.S., D.W. and B.F.H.; Methodology, Z.M., R.P., K.O.S., and B.F.H.; Software, S.V. and K.J.W., Investigation Z.M., K.J.W., K.O.S., R.H., D.W.C., R.P., D.M., K.M., R.J.E., A.N., X.L., S.X., M.B., D.M., Q.H., S.V., T.E., M.A., W.B.W., N.P., D.W., B.F.H.; Statistical analysis, Y.W., W.R.; Writing, Z.M., K.O.S., K.J.W., and B.F.H.; Funding Acquisition, B.F.H.; Resources, Y.T., C.B., N.P., D.W.; B.F.H. Supervision, B.F.H.

## STAR Methods

### LEAD CONTACT

Further information and requests for resources and reagents should be directed to and will be fulfilled by the lead contact Barton F. Haynes.

### MATERIALS AVAILABILITY

This study did not generate new unique reagents.

### DATA AND CODE AVAILABILITY

Any additional information required to reanalyze the data reported in this paper is available from the lead contact upon request.

### EXPERIMENTAL MODEL AND SUBJECT DETAILS

#### Cell line

Freestyle 293-F cell line (Thermo Fisher Scientific, Cat# R79007) was purchased from Thermo Fisher and cultured in Freestyle 293 Expression Medium (Thermo Fisher Scientific, Cat# 12338- 026). Cells were maintained in 8% CO2 at 37°C at a density between 0.3x10^6^/ml to 3x10^6^/ml.

Mycoplasma test was performed when a new stock vial was thawed at Duke University Cell Culture Facility.

#### Animals and immunizations

The DH270 UCA dKI mice has been previously described (Saunders et al., 2019). For CH848 10.17DT gp160 mRNA-LNP immunizations, 12 DH270 UCA dKI mice were randomly split into two groups (N = 6 each group) and were immunized intramuscularly (i.m.) with 20 μg of mRNA-LNP encoding CH848 10.17DT F14 gp160 and CH848 10.17DT 113C-429GCG gp160 every two weeks for three times. For CH848 10.17DT DS trimer-ferritin NP immunizations were done similarly, except that the second and the third immunizations were only one week apart. Control group mice were injected with 20 μg of empty LNP. Bleeding was performed one week after each immunization. Necropsy was performed one week after the third immunization and blood, spleen, and lymph nodes were collected. All mice were cared for in a facility accredited by the Association for Assessment and Accreditation of Laboratory Animal Care International (AAALAC). All study protocol and all veterinarian procedures were approved by the Duke University Institutional Animal Care and Use Committee (IACUC).

### METHOD DETAILS

#### Modified mRNA production

Modified mRNAs were produced by *in vitro* transcription using T7 RNA polymerase (Megascript, Ambion) on linearized plasmids encoding codon-optimized CH848 10.17DT gp160s, CH848 10.17DT SOSIP trimers, or CH848 10.17DT trimer-ferritin NPs. All the HIV-1 modified mRNA constructs used in this study and their corresponding plasmids were listed in **Table S1**. One-methylpseudouridine (m1Ψ)-5’-triphosphate (TriLink, Cat# N-1081), instead of UTP was used to produce nucleoside-modified mRNAs. Modified mRNAs contain 101 nucleotide-long polyadenylation tails for optimized expression. Modified CH848 10.17DT SOSIPv4.1 trimer and CH848 10.17DT SOSIPv5.2.8 trimer mRNAs were capped using ScriptCap m7G capping system and ScriptCap 2′-O-methyl•transferase kit (ScriptCap, CellScript) (Pardi et al., 2013). Capping of all other *in vitro* transcribed mRNAs was performed co-transcriptionally using the trinucleotide cap1 analog, CleanCap (TriLink, Cat# N-7413). All mRNAs were purified by cellulose purification, as described (Baiersdorfer et al., 2019). All mRNAs were analyzed by agarose gel electrophoresis and were stored frozen at -20 °C.

#### Nucleoside-modified mRNA-LNP production

Nucleoside-modified mRNAs were encapsulated in LNP for mouse immunizations as previously described (Jayaraman et al., 2012; Maier et al., 2013). Modified mRNAs in aqueous phase were rapidly mixed with a solution of lipids dissolved in ethanol. LNP formulation contains ionizable cationic lipid (proprietary to Acuitas)/phosphatidylcholine/cholesterol/PEG-lipid. The cationic lipid and LNP composition are described in US patent US10,221,127 (Du, 2019).

#### Nucleoside-modified mRNA transfection in 293-F cell line

293-F cells were diluted to 0.7x10^6^ cells/ml 24 h before transfection. On the next day, cells were diluted again to 1x10^6^/ml and seeded into tissue culture plates for transfection. 3 μg of mRNAs expressing gp160s were transfected into 6 ml of cells. For soluble SOSIP trimers, 30 ml of cells were transfected with 12 μg of SOSIP-expressing mRNAs and 3 μg of Furin mRNAs. Transfection volume were doubled to 60 ml for trimer-ferritin NPs. TransIT-mRNA Transfection Kit (Mirus Cat# MIR2250) was used for mRNA transfection following the manufacturer’s instructions. Transfected cells were cultured at 37°C with 8% CO2 and shaking at 120 rpm for 48 h (for gp160) or 72 h (for SOSIP trimers and trimer-ferritin NPs) before harvest.

#### Evaluation of expression and folding of modified mRNA-expressed CH848 10.17DT gp160s, SOSIP trimers, and trimer-ferritin NPs

The expression and folding of modified mRNA-encoded CH848 10.17DT transmembrane gp160s, Soluble SOSIP trimers, and trimer-ferritin NPs were defined as follows. For CH848 10.17DT transmembrane gp160s, flow cytometry was used to measure binding of a panel of bnAbs and nnAbs. BnAb binding reactivity indicated successful expression of gp160 Envs on cell surface with desired antigenicity. Binding of nnAbs 17b and 19b measured the ability of various stabilizing mutations to keep the gp160 Envs in prefusion conformation and to decrease the exposure of non-neutralizing epitopes CCR5 binding site and distal V3 loop. Additionally, 7B2 binding was used to measure the exposure of gp41.

For CH848 10.17DT soluble trimer, total Env forms were purified by *Galanthus nivalis* lectin (GNL) from modified mRNA-transfected 293-F cell supernatant. A panel of nnAbs and bnAbs were used in ELISA to measure the expression of non-neutralizing and neutralizing epitopes. Binding of nnAbs 17b and 19b after CD4 triggering were measured by SPR. To assess the percent of trimeric Envs in GNL-purified materials, size-exclusion ultra-performance liquid chromatography (SE-UPLC) analysis was performed with PGT151 affinity-purified CH848 10.17DT SOSIP trimer as a standard. Finally, Negative-stain Electron Microscopy (NSEM) analysis was performed to confirm trimer formation in GNL-purified 293-F transfection supernatant.

Similar antigenicity measurement and SPR analysis was performed on GNL-purified 293-F cell supernatant transfected with modified mRNA expressing CH848 10.17DT trimer-ferritin NPs to evaluate the expression of well-folded Env trimers on the ferritin nanoparticle. Additionally, transfected 293-F supernatant were affinity purified with PGT145-conjugated beads, which exclude host glycan proteins that may be purified by GNL. PGT145-purified materials were analyzed by NSEM to confirm the assembly of CH848 10.17DT DS ferritin NPs.

##### Flow cytometry

Binding of bnAbs to CH848 10.17DT gp160s was performed by flow cytometry as previously described (Henderson et al., 2020; Saunders et al., 2021). Briefly, modified mRNA-transfected 293-F cells were harvested 48 h after transfection and were washed once with 1% BSA in PBS. Then, cells were incubated with 10 μg/ml of bnAbs in V-bottom 96-well plates for 30 min at 4 °C. Cells were then washed with 1% BSA in PBS and incubated with Goat F(ab’)2 Anti-Human IgG - (Fab’)2 (PE) (Abcam Cat# ab98606, RRID:AB_10672217) for 30 min at 4 °C in dark. Then, cells were washed once with PBS and dead cells were stained with LIVE/DEAD Fixable Aqua Dead Cell Stain Kit (Invitrogen Cat# L34957, 1:1000 dilution in PBS) for 15 min at 4 °C in dark, then washed twice and re-suspended in 1% BSA in PBS. Flow cytometric data were acquired on a LSRII High-throughput system using FACSDIVA software (BD Biosciences) and were analyzed with FlowJo software (FlowJo). The percentage of 293-F cells that were PE positive was shown in the results.

Measurement of binding of nnAbs after CD4 treatment has been described previously (Henderson et al., 2020). Briefly, mRNA-transfected 293-F cells were first incubated with 20 μg/ml of soluble CD4 (sCD4), eCD4-Ig or CD4-IgG2 for 10 min at 4 °C. Cells were washed once with 1% BSA in PBS and then incubated with 10 μg/ml of nnAbs 17b, 19b or 7B2 for 30 min at 4°C. Then, cells were incubated with Goat F(ab’)2 Anti-Human IgG - (Fab’)2 (PE) and dead cells were stained with LIVE/DEAD Fixable Aqua Dead Cell Stain Kit. Data acquisition and analysis were the same as described above.

##### Galanthus nivalis lectin purification of SOSIP trimers and trimer-ferritin NPs

293-F cells transfected with modified mRNAs expressing SOSIP trimers or trimer-ferritin NPs were harvested 72 h after transfection and were centrifuged for 30 min at 3000 rpm to remove cells and debris. Supernatant were first filtered using a 0.22 μm vacuum filter and were then concentrated by 50-fold using 10 kDa MWCO concentrators. Concentrated supernatant was incubated with 200 μl of agarose bound *Galanthus nivalis* lectin (GNL) (Vector Laboratories Cat# AL-1243) with gentle rotation at 4 °C overnight. The next day, GNL agarose beads were washed with MES wash buffer (20 mM MES, 130 mM NaCl, 10 mM CaCl2 pH 7.0) for three times, and SOSIP trimers or trimer-ferritin NPs were eluted by 500 mM Methyl alpha-D-mannopyranoside in MES wash buffer. The eluates were then dialyzed to 10 mM Tris-HCl pH8 500 mM NaCl using 30kDa MWCO spin concentrators. GNL-purified SOSIP trimers or trimer-ferritin NPs were snap- freezed and stored in -80 °C.

##### Enzyme-linked immunosorbent assay (ELISA)

Binding reactivity of modified mRNA-expressed CH848 10.17DT SOSIP trimers and trimer-ferritin NPs to bnAbs and nnAbs was measured by enzyme-linked immunosorbent assay (ELISA). In brief, HIV-1 antibodies were coated onto 384-well assay plates in 0.1M Sodium bicarbonate overnight at 4 °C. GNL-purified SOSIP trimers or trimer-ferritin NPs with serial dilutions were then captured on the plates. Next, poly-serum from CH848 10.17DT-immunized rhesus macaque was incubated for 1 h at room temperature. Then, Mouse Anti-Monkey IgG-HRP (SouthernBiotech Cat# 4700-05, RRID:AB_2796069) was incubated for 1 h at room temperature and plates were developed with SureBlue Reserve TMB 1-Component Microwell Peroxidase Substrate (Seracare Cat# 5120-0083) for 15 min and were stopped with 1% HCl solution. Absorbance at 450 nm were determined by SpectraMax Plus 384 microplate reader (Molecular Devices) and log area-under-curve (log AUC) were calculated using Prism (Graphpad) and shown in figures.

The base binding antibody assay was performed similarly. Briefly, base binding antibody DH1029 was coated onto plates to capture samples. Then, a rabbit serum was incubated before detection with Goat polyclonal Secondary Antibody to Rabbit IgG - H&L (HRP) (Abcam Cat# ab97080, RRID:AB_10679808). The plate development, data acquisition, and analysis were the same as described above.

##### Size-exclusion ultra-performance liquid chromatography (SE-UPLC)

Size exclusion chromatography of modified mRNA-expressed GNL-purified CH848 10.17DT SOSIP trimers was performed using a Waters Acquity H-Class Bio UPLC System with a Waters Acquity UPLC BEH SEC 450Å, 2.5 µm, 4.6 x 150 mm column (Waters Corporation). An isocratic elution with a mobile phase of 20 mM sodium phosphate 300 mM NaCl pH 7.4, and a flow rate of 0.2 ml/min, was used for the analysis with a quaternary pump. Samples and protein standards were maintained at 5-8°C in the auto-sampler rack prior to injection at a volume of 10 µl. Samples and protein standards with a concentration greater than 1.0 mg/ml were diluted to a down to 1.0 mg/mL using Type 1 water. The column temperature was set to 30 °C with detection at a wavelength of 214 nm using a photodiode array detector.

##### Negative-stain Electron Microscopy (NSEM)

Negative-stain electron microscopy (NESM) analysis of modified mRNA-expressed CH848 10.17DT SOSIP trimers and trimer-ferritin NPs were performed as previously described (Saunders et al., 2017; Williams et al., 2021).

##### Surface Plasmon Resonance (SPR)

SPR analyses of modified mRNA-expressed SOSIP proteins incubated with and without sCD4 against distal V3 loop antibody 19b and CCR5 binding site antibody 17b were obtained using the Biacore S200 instrument (Cytiva). Antibodies 19b and 17b were immobilized onto a CM3 sensor chip to a level of 2000-4000RU. A negative control Influenza IgG1 antibody (CH65) was also immobilized onto the sensor chip for reference subtraction. Modified mRNA-expressed GNL-purified CH848 10.17DT SOSIP trimers or trimer-ferritin NPs were diluted down in HBS-N 1x running buffer to 0.5-2.0 μg and incubated with a 2-8x higher dose of soluble CD4 (4.4 μg) (Progenics Therapeutics). Proteins incubated with and without sCD4 were injected over the sensor chip surface using the High performance injection type for 180s at 30 μl/min. The protein was then allowed to dissociate for 600s followed by sensor surface regeneration of two 20 s injections of glycine pH 2.0 at a flow rate of 50 μl/min. Results were analyzed using the BIAevaluation Software (Cytiva). Protein binding to the CH65 immobilized sensor surface as well as buffer binding were used for double reference subtraction to account for non-specific protein binding and signal drift.

#### Mouse serological analysis by ELISA

##### Serum IgG antigen binding assay

Serum IgG binding to HIV-1 antigens was measured by ELISA as previously described (Saunders et al., 2019).

##### DH1029 blocking assay

DH1029 blocking by vaccinated mouse sera was performed in ELISA. Briefly, 384-well assay plate were coated with 2 μg/ml PGT145. Then, 0.125 μg/ml of CH848 10.17DT SOSIP trimer were captured for 1 h at room temperature. Next, mouse sera at 1:50 dilution or DH1029 mAb in serial dilution were incubated for 1 h. Next, biotinylated DH1029 were added to the plate for 1 h and binding were detected by High Sensitivity Streptavidin-HRP (Thermo Fisher Scientific, Cat #21130). Plate development and data acquisition were the same as described above.

##### HIV-1 pseudovirus neutralization assay

Neutralization assays were performed in TZM-bl reporter cells as described (Mascola et al., 2005).

#### Next-generation sequencing (NGS)

We performed next-generation sequencing (NGS) on mouse antibody heavy and light chain variable genes using an Illumina sequencing platform. First, RNA was purified from splenocytes using a RNeasy Mini Kit (Qiagen, Cat# 74104). Purified RNA was quantified via Nanodrop (Thermo Fisher Scientific) and used to generate Illumina-ready heavy and light chain sequencing libraries using the SMARTer Mouse BCR IgG H/K/L Profiling Kit (Takara, Cat# 634422). Briefly, 1 μg of total purified RNA from splenocytes was used for reverse transcription with Poly dT provided in the SMARTer Mouse BCR kit for cDNA synthesis. Heavy and light chain genes were then separately amplified using a 5’ RACE approach with reverse primers that anneal in the mouse IgG constant region for heavy chain genes and IgK for the light chain genes (SMARTer Mouse BCR IgG H/K/L Profiling Kit). The DH270 UCA KI mouse model has the light chain gene knocked into the kappa locus, therefore kappa primers provided in the SMARTer Mouse BCR kit were used for light chain gene library preparation. 5 µl of cDNA was used for heavy and light chain gene amplification via two rounds of PCR; PCR1 used 18 cycles and PCR2 used 12 cycles. During PCR2, Illumina adapters and indexes were added. Illumina-ready sequencing libraries were then purified and size-selected by AMPure XP (Beckman Coulter, Cat# A63881) using kit recommendations. The heavy and light chain libraries per mouse were indexed separately, thus allowing us to deconvolute the mouse-specific sequences during analysis. Libraries were quantified using QuBit Fluorometer (Thermo Fisher). Mice were pooled by groups for sequencing on the Illumina MiSeq Reagent Kit v3 (600 cycle) (Illumina, Cat# MS-102-3003) using read lengths of 301/301 with 20% PhiX.

#### Antibody sequence analysis

NGS data analysis and the analysis of improbable mutation frequencies was performed as described (Wiehe et al., 2018).

#### Flow cytometric phenotyping of GC responses

For immunophenotyping of murine B cells and Tfh cells, spleens from immunized mice one week after the third immunization were processed into single-cell suspensions and treated with ACK lysis buffer to remove red blood cells. Splenocytes (2x10^6^) were suspended in 100 μL PBS/2% FBS. To detect antigen-specific B cells, fluorochrome-mAb conjugates and fluorochrome-conjugated CH848 10.17DT Envs were prepared as a master mix at 2x concentration, then 100 μL of 2x master mix was added to an equal volume of cells (**Figure S4**). Staining for T cell subsets was conducted in the same manner, with the additional step for detection of biotinylated mAb with Streptavidin–APC. Cells were incubated at 4°C for 20 minutes, then washed with PBS. Cells were resuspended in 100 μL PBS containing Near-IR Live/Dead (Thermo Fisher Scientific) at 1:1000, and incubated at room temperature for 20 min. Cells were washed in PBS/2% FBS, then re- suspended in PBS/2% formaldehyde. Cells were analyzed on a BD LSRII (BD Biosciences). Data were analyzed using FlowJo v10 (FlowJo).

#### Isolation of CH848 10.17DT-specific neutralizing monoclonal antibodies (mAbs)

##### Antibody cloning, screening, and mAbs expression

Immunoglobulin (Ig) gene were cloned from sorted single B cells as previously described (Liao et al., 2009). Briefly, complementary DNA (cDNA) of Ig genes were amplified by reverse-transcription with SuperScript III First-Strand Synthesis System (Thermo Fisher Scientific, Cat# 18080051) using random hexamer oligonucleotides as primers. Ig gene cDNA was then used as template in nested PCR for heavy and light chain gene amplification using AmpliTaq Gold 360 Master Mix (Thermo Fisher Scientific, Cat #4398881). Mouse Ig-specific primers and DH270 variable region-specific primers were used to amplify mouse endogenous Ig genes and DH270 KI Ig genes. Agarose gel electrophoresis was used to identify positive PCR amplification and Ig genes were recovered by Sanger sequencing. Following sequencing, contigs of PCR amplicon sequences were assembled, and Ig genes were inferred with human Ig gene library and mouse Ig gene library in Cloanalyst. PCR reactions with successful Ig sequence recovery were purified using AMPure XP kit (Beckman Coulter, Cat# A63881). Purified PCR product was used for overlapping PCR to generate a linear antibody expression cassette. The expression cassette was transiently transfected with into 293i cells with ExpiFectamine 293 Transfection Kit (Thermo Fisher Scientific, Cat# A14525). The supernatant was harvested 72 h after transfection and screened in ELISA binding assays with a panel of protein of interests. The genes of selected heavy chains were synthesized with human IgG1 backbone (GenScript). Kappa and lambda chains were synthesized similarly. To express mAbs plasmids were prepared for transient transfection using the Plasmid Plus Mega Kit (Qiagen, Cat #12981). Heavy and light chain plasmids were co-transfected into 293i cells using ExpiFectamine 293 Transfection Kit for antibody production.

### QUANTIFICATION AND STATISTICAL ANALYSIS

Exact Wilcoxon Mann-Whitney U tests were performed without any adjustment for multiple comparisons. Significant results were indicated in figures and figure legends as: * P < 0.05; ** P< 0.01.

Table S3. CH848 10.17DT trimer-ferritin NP mRNA-LNP vaccine-induced monoclonal antibodies ELISA binding magnitudes. Related to Figure 7.

