## Supplemental figures for "Ability of nucleoside-modified mRNA to encode HIV-1 envelope trimer nanoparticles"

### SUPPLEMENTAL INFORMATION

**Table S1.** Stabilizing mutations studied for modified mRNA-encoded CH848 10.17DT transmembrane gp160s, soluble SOSIP trimers, and trimer-ferritin nanoparticles. **Related to Figure 1.**

| Env form | Design | Mutations | Reference |
|---|---|---|---|
| Transmembrane gp160s | DS | 201C-433C | Kwon et al., 2015 |
| | F14 | 68I, 204V, 208L, 255L | Henderson et al., 2020 |
| | Vt8 | 203M, 300L, 302L, 320M, 422M | Henderson et al., 2020 |
| | F14/Vt8 | F14+Vt8 | Henderson et al., 2020 |
| | 113C-429GCG | 113C-429C, 428G, 430G | Zhang et al., 2018 |
| | 113C-431GCG | 113C-431C, 430G, 432G | Zhang et al., 2018 |
| Soluble SOSIP trimers | v4.1 | 501C-605C, 559P, R6, ΔMPER, 535M, 543N/Q, 316W, 64K | De Taeye et al., 2015 |
| | v5.2.8 | V4.1, 66R, 73C-561C, L165, Q432, R429, K65, T106, E49, D47, R500 | Guenaga et al., 2015 |
| | DS | 201C-433C | Kwon et al., 2015 |
| | F14 | 68I, 204V, 208L, 255L | Henderson et al., 2020 |
| | Vt8 | 203M, 300L, 302L, 320M, 422M | Henderson et al., 2020 |
| | F14/Vt8 | F14+Vt8 | Henderson et al., 2020 |
| | UFO | HR-1 re-designed, 501C-605C | Kong et al., 2016 |
| | v5.2.8+UFO | v5.2.8+UFO | Guenaga et al., 2015; Kong et al., 2016 |
| Multimeric trimer-ferritin NPs | DS | 201C-433C | Kwon et al., 2015 |
| | 113C-429GCG | 113C-429C, 428G, 430G | Zhang et al., 2018 |

**Table S2.** Plasmids used for *in vitro* transcription of each mRNA construct. **Related to Figure 1.**

| **mRNA Construct** | **Plasmid** |
|---|---|
| CH848 10.17DT gp160 | ccap1-TEV-CH848-10.17-DTgp160-A101 |
| CH848 10.17DT DS gp160 | ccap1-TEV-CH848-10.17-DT-DS gp160-A101 |
| CH848 10.17DT F14 gp160 | ccap1-TEV-CH0848-10.17-DTgp160_F14-A101 |
| CH848 10.17DT Vt8 gp160 | ccap1-TEV-CH848-10.17-DTgp160-VT8-A101 |
| CH848 10.17DT F14/Vt8 gp160 | ccap1-TEV-CH0848-10.17-DTgp160_F14_VT8-A101 |
| CH848 10.17DT 113C-429GCG gp160 | ccap1-TEV-CH0848.d949.10.17gp160_N133DN138T_113C-429GCG-A101 |
| CH848 10.17DT 113C-431GCG gp160 | ccap1-TEV-CH0848.d949.10.17gp160_N133DN138T_113C-431GCG-A101 |
| CH848 10.17DT SOSIPv4.1 trimer | cap1-TEV-CH848.3.D0949.10.17CHIM1.6R.SOSIP.664V4.1_N133D N138T-A101 |
| CH848 10.17DT SOSIPv5.2.8 trimer | cap1-TEV-CH848.3.D0949.10.17CHIM.SOSIPV5.2.8_N133DN138T-A101 |
| CH848 10.17DT F14 SOSIP trimer | ccTEV-CH848-10.17-DT-F14 SOSIP-A101 |
| CH848 10.17DT Vt8 SOSIP trimer | ccTEV-CH848-10.17-DT-Vt8 SOSIP-A101 |
| CH848 10.17DT F14/Vt8 SOSIP trimer | ccTEV-CH848-10.17-DT-F14Vt8 SOSIP-A101 |
| CH848 10.17DT 113C-429GCG SOSIP trimer | ccap1-TEV-CH848.3.10.17ch.SOSIP_DT_113C-429GCG-A101 |
| CH848 10.17DT UFO-BG SOSIP trimer | ccap1-TEV-CH848.d949.10.17 N133D N138T SOSIP.UFO-BG-A101 |
| CH848 10.17DT DS trimer-ferritin NP | ccap1-TEV-CH848.3.D0949.10.17chDS.SOSIP_N133DN138T-ferritin-A101 |
| CH848 10.17DT 113C-429GCG trimer-ferritin NP | ccap1-TEV-CH848.10.17ch.SOSIP_DT_113C-429GCG-ferritin-A101 |
| CH848 10.17DTe DS 3xGS linker trimer-ferritin NP | CH848.3.D0949.10.17_N113D_N138T_D230N_H289N_P291S_E169K_DS_SOSIP_LSK_ferritin_3xGSlinker |
| CH848 10.17DTe DS 2xGS linker trimer-ferritin NP | CH848.3.D0949.10.17_N113D_N138T_D230N_H289N_P291S_E169K_DS_SOSIP_LSK_ferritin_2xGSlinker |
| CH848 10.17DTe DS VRC trimer-ferritin NP | CH848.3.D0949.10.17_N133D_N138T_D230N_H289N_P291S_E169K_DS.SOSIP_VRCferritin |

**A**

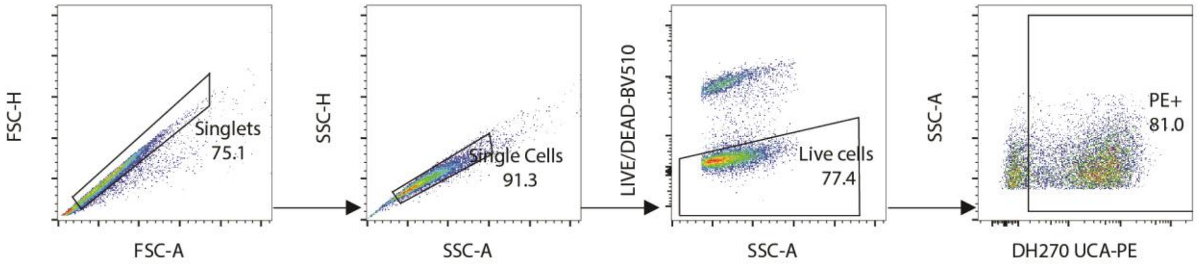


**B**

| | CH65 | CH01 | PG9 | PGT125 | DH270 UCA |
|---|---|---|---|---|---|
| **10.17DT** | 0.07 | 37.30 | 9.87 | 82.90 | 76.07 |
| **10.17DT DS** | 0.12 | 25.33 | 15.61 | 64.97 | 39.60 |
| **10.17DT F14** | 0.07 | 37.57 | 6.45 | 85.73 | 81.63 |
| **10.17DT Vt8** | 0.09 | 13.31 | 1.65 | 86.13 | 34.77 |
| **10.17DT F14/Vt8** | 0.09 | 8.48 | 1.15 | 82.50 | 35.40 |
| **10.17DT 113C-429GCG** | 0.07 | 35.27 | 0.89 | 86.37 | 78.03 |
| **10.17DT 113C-431GCG** | 0.04 | 18.33 | 0.90 | 85.87 | 77.07 |

**C**

| | CH65 | 17b | 19b | 7B2 |
|---|---|---|---|---|
| **10.17DT** | | | | |
| No CD4 | 0.07 | 4.77 | 3.32 | 6.63 |
| +sCD4 | 0.07 | 43.93 | 26.77 | 8.29 |
| +eCD4-Ig | 0.07 | 72.77 | 69.83 | 26.67 |
| +CD4-IgG2 | 0.13 | 58.40 | 54.07 | 8.01 |
| **10.17DT DS** | | | | |
| No CD4 | 0.36 | 0.68 | 13.68 | 21.53 |
| +sCD4 | 0.41 | 0.53 | 14.73 | 20.83 |
| +eCD4-Ig | 0.38 | 0.37 | 17.84 | 20.60 |
| +CD4-IgG2 | 0.39 | 0.53 | 22.74 | 19.60 |
| **10.17DT F14** | | | | |
| No CD4 | 0.07 | 1.61 | 1.86 | 6.53 |
| +sCD4 | 0.04 | 1.95 | 2.06 | 6.17 |
| +eCD4-Ig | 0.05 | 3.86 | 4.19 | 5.44 |
| +CD4-IgG2 | 0.04 | 2.48 | 2.60 | 6.16 |
| **10.17DT Vt8** | | | | |
| No CD4 | 0.70 | 0.86 | 2.41 | 11.52 |
| +sCD4 | 0.54 | 0.87 | 4.24 | 11.72 |
| +eCD4-Ig | 0.58 | 11.76 | 14.60 | 10.52 |
| +CD4-IgG2 | 0.60 | 11.78 | 13.76 | 11.29 |
| **10.17DT F14/Vt8** | | | | |
| No CD4 | 0.78 | 1.13 | 7.63 | 5.62 |
| +sCD4 | 0.24 | 0.40 | 7.11 | 6.46 |
| +eCD4-Ig | 0.24 | 0.53 | 9.56 | 5.70 |
| +CD4-IgG2 | 0.30 | 0.49 | 14.20 | 6.06 |
| **10.17DT 113C-429GCG** | | | | |
| No CD4 | 0.07 | 1.88 | 2.26 | 7.65 |
| +sCD4 | 0.04 | 2.03 | 2.33 | 8.37 |
| +eCD4-Ig | 0.08 | 1.74 | 2.25 | 8.45 |
| +CD4-IgG2 | 0.06 | 2.11 | 2.23 | 10.10 |
| **10.17DT 113C-431GCG** | | | | |
| No CD4 | 0.04 | 2.62 | 2.53 | 10.09 |
| +sCD4 | 0.07 | 2.67 | 2.50 | 9.59 |
| +eCD4-Ig | 0.09 | 2.52 | 2.40 | 6.32 |
| +CD4-IgG2 | 0.14 | 2.43 | 2.49 | 10.02 |

**Figure S1. Antigenicity of modified mRNA-encoded CH848 10.17DT transmembrane gp160s. Related to Figure 1.**

**(A)** Gating strategy for flow cytometric analysis of antibody binding to modified mRNA-expressed CH848 10.17DT gp160s on 293-F cell surface. Figure shows representative gating of DH270 UCA binding to CH848 10.17DT gp160.

**(B)** Binding of bnAbs to modified mRNA-expressed CH848 10.17DT transmembrane gp160s. Data shown were means of PE+ cell percentage among total live cells from three independent experiments. Influenza hemagglutinin Ab CH65 was used as negative control.

**(C)** Binding of nnAbs 17b, 19b, and 7B2 to CH848 10.17DT gp160 after CD4 triggering. Three forms of CD4: soluble CD4 (sCD4), eCD4-Ig, and CD4-IgG2 were used. Binding of Abs were detected by Goat anti-Human IgG F(ab')2 secondary antibody labeled with PE. Data shown were means of PE+ cell percentage among total live cells from three independent experiment. Influenza hemagglutinin Ab CH65 was used as negative control.

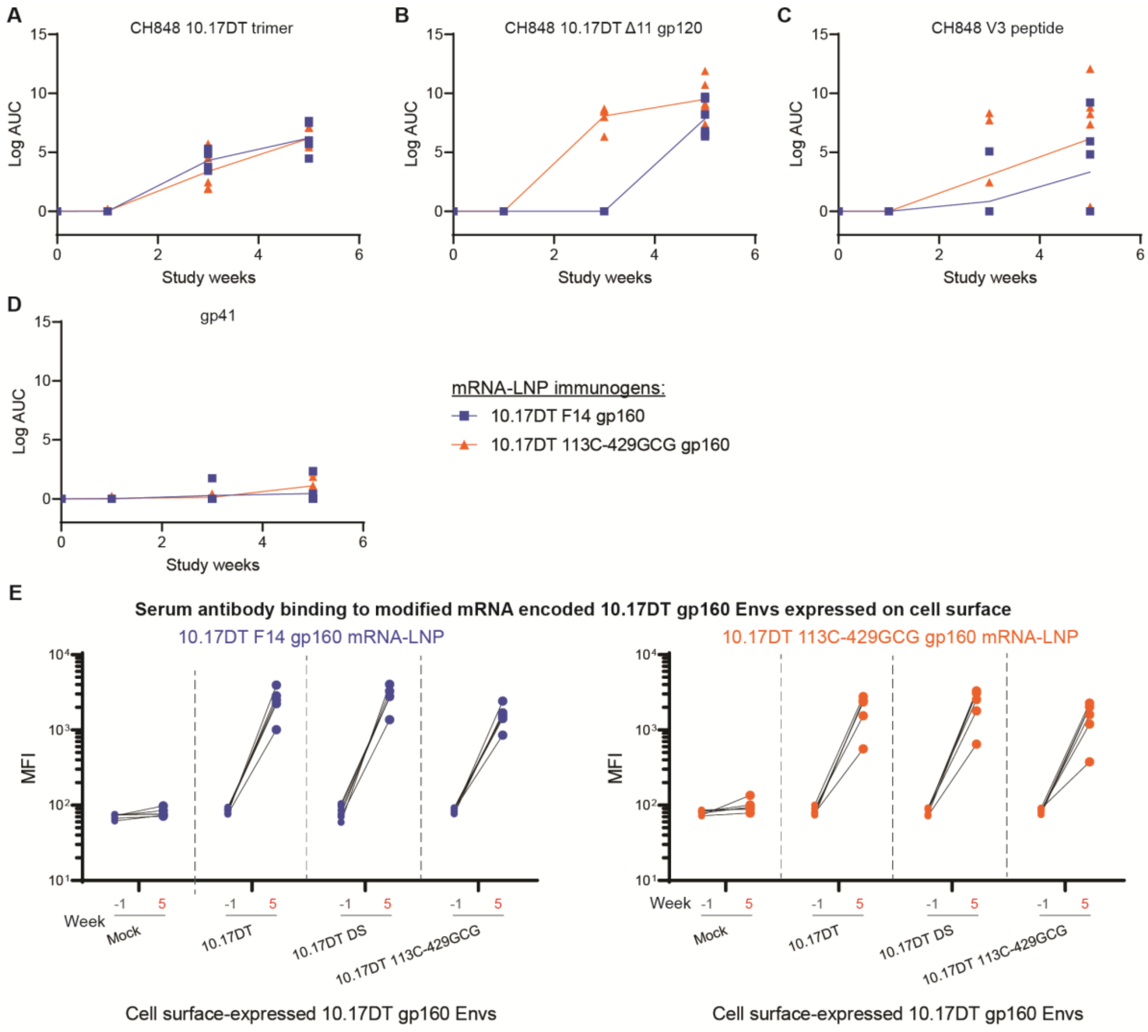


**F**

| Immunization | IC50 (1/dilution) in TZM-bl cells | | | | | | | |
|---|---|---|---|---|---|---|---|---|
| | MuLV | 92RW020 | CH848.d949.10.17 | 6101 | CH848.d949.10.17.N332T | Q23 | CH848.d949.10.17.DT | CH848.d949.10.17.DT.D230N.H289N.P291S |
| CH848 10.17DT F14 gp160 mRNA-LNP | <200 | <200 | <200 | <200 | <200 | <200 | 4586 | 4016 |
| | <200 | <200 | 204 | <200 | <200 | <200 | 8868 | 6898 |
| | <200 | <200 | <200 | <200 | <200 | <200 | 8623 | 6797 |
| | <200 | <200 | 406 | <200 | <200 | <200 | 38384 | 24220 |
| | <200 | <200 | 584 | <200 | <200 | <200 | 16195 | 11164 |
| | <200 | <200 | 745 | <200 | <200 | <200 | 23994 | 21662 |
| CH848 10.17DT 113C-429GCG gp160 mRNA-LNP | <200 | <200 | 615 | <200 | <200 | <200 | 16862 | 13945 |
| | <200 | <200 | 424 | <200 | <200 | <200 | 6709 | 7092 |
| | <200 | <200 | 480 | <200 | <200 | <200 | 5896 | 5534 |
| | <200 | <200 | 331 | <200 | <200 | <200 | 7800 | 4540 |
| | <200 | <200 | <200 | <200 | <200 | <200 | 8501 | 7959 |
| | <200 | <200 | <200 | <200 | <200 | <200 | 36295 | 16434 |

| ID50 |
|---|
| nt |
| <200 |
| 160-999 |
| 1000-9999 |
| >10000 |

**Figure S2. Serum antibody binding and neutralization activity after CH848 10.17DT gp160 mRNA-LNP immunizations in DH270 UCA KI mice. Related to Figure 2.**

**(A-D)** ELISA logAUCs of vaccinated mouse serum IgG binding to **(A)** CH848 10.17DT DS SOSIP trimer; **(B)** CH848 10.17 Δ11 gp120; **(C)** CH848 V3 loop peptide; **(D)** gp41 subunit.

**(E)** Week 5 serum antibody binding to modified mRNA-expressed CH848 10.17DT, CH848 10.17DT DS, and CH848 10.17DT 113C-429GCG gp160s on 293-F cell surface. Each dot signifies an individual mouse (N = 6 each group). Data shown are MFI measured by flow cytometry.

**(F)** Serum neutralizing titers after CH848 10.17DT gp160 mRNA-LNPs immunization. Serum neutralization titers one week after the third immunization in DH270 UCA KI mice were measured using TZM-bl reporter cells. Data shown were ID50s. Murine leukemia virus (MuLV) was used as negative control.

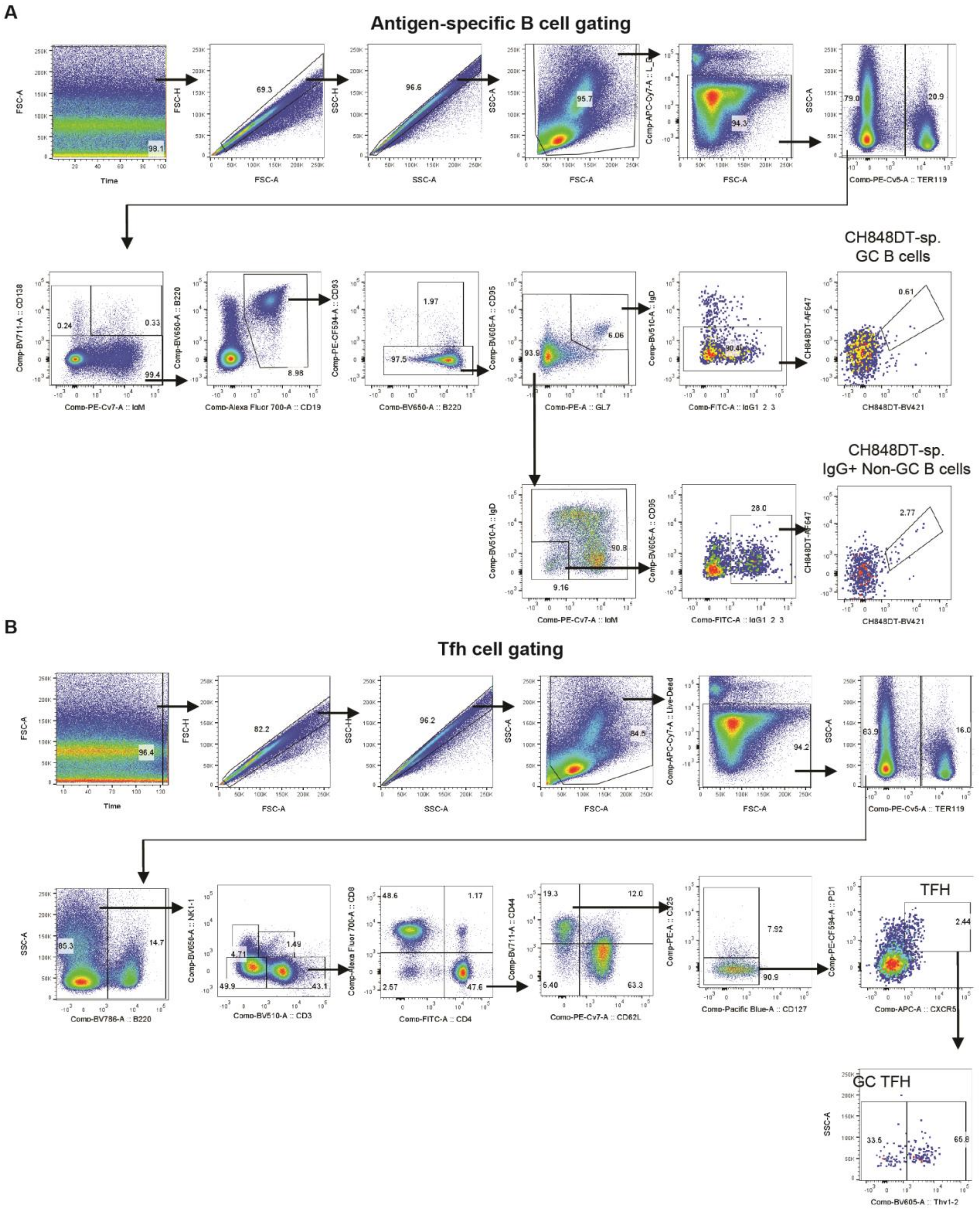


**Figure S3. Flow cytometric gating strategies for antigen-specific B cells and Tfh cell populations. Related to Figures 2 and 5.**

**(A**) The gating strategy to identify CH848 10.17DT-specific germinal center B cells and memory B cells.

**(B)** The gating strategy to identify T follicular helper (Tfh) and germinal center Tfh (GC Tfh) cells.

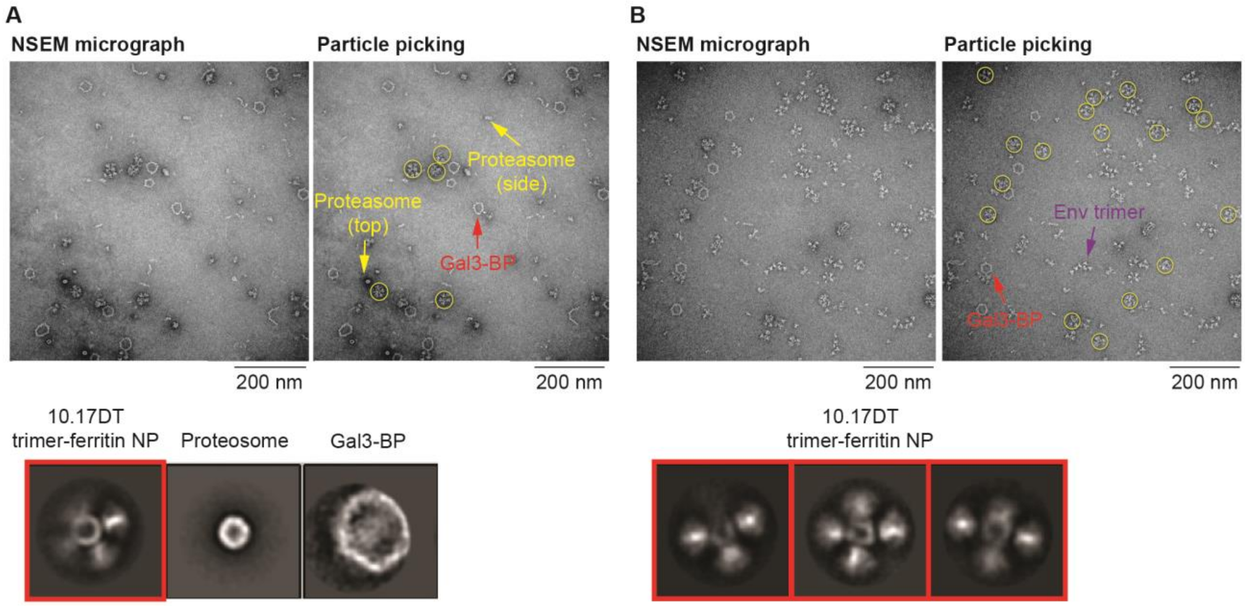


**Figure S4. Modified mRNA-expressed CH848 10.17DT trimer-ferritin fusion protein assembles into well-folded NPs. Related to Figure 3.**

**(A, B)** NSEM analysis of modified mRNA-expressed, PGT145-purified **(A)** CH848 10.17DT DS VRC trimer-ferritin NPs or **(B)** CH848 10.17DT DS 3xGS linker trimer-ferritin NPs. Representative NSEM micrograph is shown on top left, and particle picking is shown on top right. Below are 2D class averages of each sample. Yellow circle, trimer-ferritin NP; purple arrow, free Env trimer; yellow arrow, proteasome; red arrow, Galectin-3 binding protein (Gal3-BP). Scale bar, 200 nm.

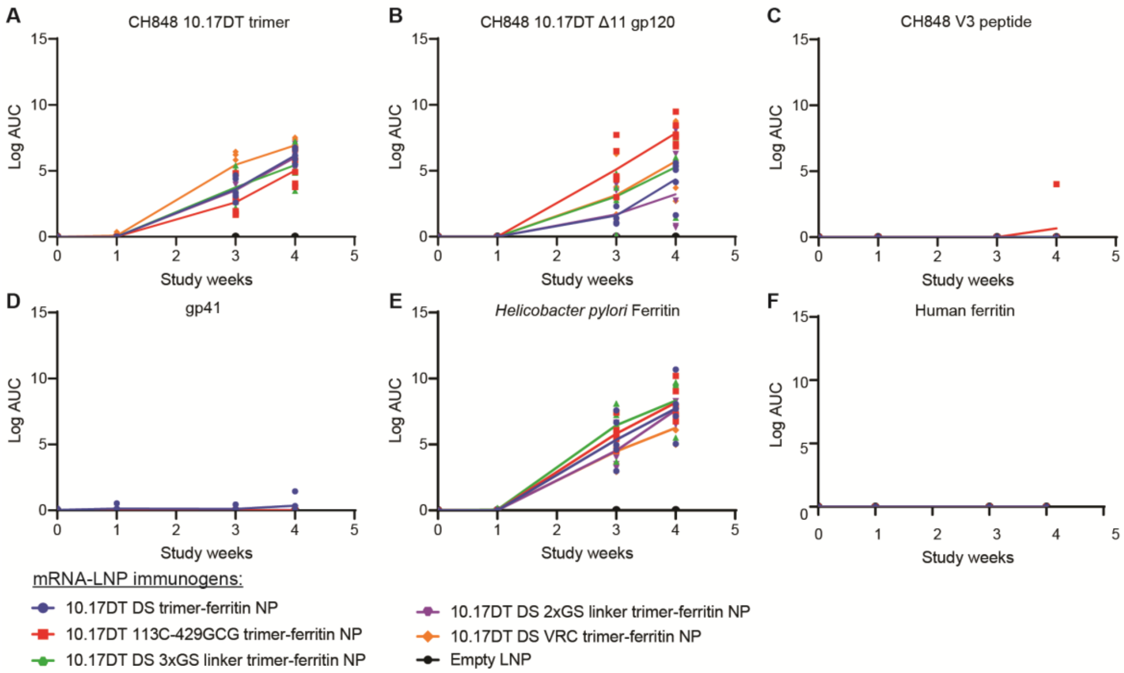


**G**

| | ID50 (dilution) in TZM-bl cells | | | | | | | |
|---|---|---|---|---|---|---|---|---|
| | MuLV | CH848.10.17 | CH848.10.17.N332T | CH848.10.17.DT | CH848.10.17.DT.N332T | CH848.10.17.DT.D230N.H289N.P291S | 92RW020.2 | 6101.10 |
| Immunization | | | | | | | | |
| CH848 10.17DT DS ferritin NP mRNA-LNP | <160 | <160 | nt | 4344 | <160 | 5301 | <160 | 170 |
| | <160 | <160 | nt | 7134 | <160 | 5542 | <160 | <160 |
| | <160 | <160 | nt | 6034 | <160 | 6018 | 172 | 285 |
| | <160 | <160 | nt | 4682 | <160 | 3480 | <160 | 185 |
| | <160 | 194.5 | <160 | 7362 | <160 | 7803 | <160 | <160 |
| CH848 10.17DT 113C-429GCG ferritin NP mRNA-LNP | <160 | <160 | nt | 4187 | <160 | 5321 | <160 | 197 |
| | <160 | <160 | nt | 1966 | <160 | 1629 | <160 | <160 |
| | <160 | 532 | <160 | 15206 | <160 | 20932 | <160 | 177 |
| | <160 | <160 | nt | 4071 | <160 | 5360 | <160 | 168 |
| | <160 | 532 | <160 | 4396 | <160 | 6340 | <160 | 172 |
| | <160 | <160 | <160 | 1519 | <160 | 1936 | <160 | 318 |
| CH848 10.17DT DS 3xGS linker LSK ferritin NP mRNA-LNP | <160 | <160 | nt | 12110 | <160 | 14835 | <160 | 186 |
| | <160 | <160 | nt | 4549 | <160 | 6242 | <160 | 192 |
| | <160 | <160 | nt | 12339 | <160 | 23328 | <160 | 190 |
| | <160 | <160 | nt | 6308 | <160 | 7913 | <160 | <160 |
| | <160 | <160 | nt | 3590 | <160 | 5466 | <160 | 219 |
| | <160 | <160 | nt | 1501 | <160 | 1187 | <160 | <160 |
| CH848 10.17DT DS 2xGS linker LSK ferritin NP mRNA-LNP | <160 | 324.5 | <160 | 10127 | <160 | 16398 | <160 | <160 |
| | <160 | <160 | nt | 5041 | <160 | 6892 | <160 | <160 |
| | <160 | <160 | nt | 6573 | <160 | 12384 | <160 | 185 |
| | <160 | <160 | nt | 4749 | <160 | 5580 | <160 | 163 |
| | <160 | <160 | nt | 5448 | <160 | 9133 | <160 | 225 |
| CH848 10.17DT DS VRC ferritin NP mRNA-LNP | <160 | <160 | nt | 9814 | <160 | 15247 | <160 | 160 |
| | <160 | 415.5 | <160 | 13412 | <160 | 24893 | <160 | 360 |
| | <160 | 325 | <160 | 14835 | <160 | 18650 | <160 | 492 |
| | <160 | 467 | <160 | 20052 | <160 | 36649 | <160 | 193 |
| | <160 | <160 | nt | 3117 | <160 | 7575 | <160 | 172 |
| | <160 | <160 | nt | 4399 | <160 | 8302 | <160 | 261 |
| Empty LNP | <160 | <160 | nt | <160 | <160 | <160 | <160 | <160 |
| | <160 | <160 | nt | <160 | <160 | <160 | <160 | <160 |
| | <160 | <160 | nt | <160 | <160 | <160 | <160 | 261 |
| | <160 | <160 | nt | <160 | <160 | <160 | <160 | 432 |
| | <160 | <160 | nt | <160 | <160 | <160 | <160 | 225 |
| | <160 | <160 | nt | <160 | <160 | <160 | <160 | <160 |

| ID50 |
|---|
| nt |
| <160 |
| 160-999 |
| 1000-9999 |
| >10000 |

**Figure S5. Serum IgG binding activity after CH848 10.17DT trimer-ferritin NP mRNA-LNP immunizations in DH270 UCA dKI mice. Related to Figure 5.**

**(A-F)** ELISA logAUCs of vaccinated mouse serum IgG binding to **(A)** CH848 10.17DT DS SOSIP trimer; **(B)** CH848 10.17 Δ11 gp120; **(C)** CH848 V3 loop peptide; **(D)** gp41 subunit; **(E)** *Helicobacter pylori* ferritin**,** and **(F)** human ferritin.

**(G)** Serum neutralizing titers after CH848 10.17DT trimer-ferritin NP mRNA-LNP immunizations. Serum neutralization titers one week after the third immunization in DH270 UCA KI mice were measured using TZM-bl reporter cells. Data shown were ID50s. Murine leukemia virus (MuLV) was used as negative control.

A

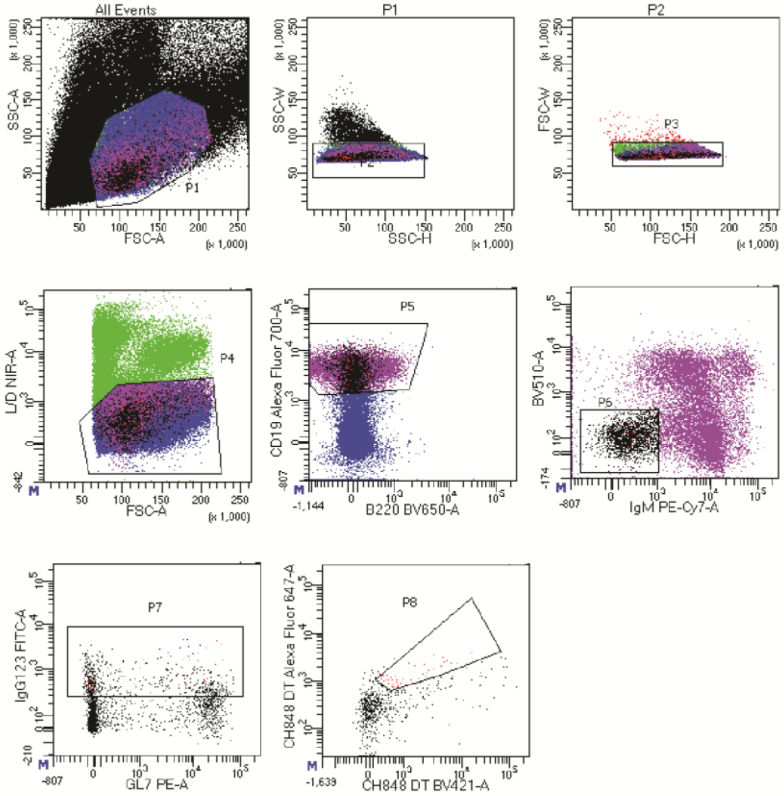


B

| Group | Animal ID | mAb # | VH1-2 # | | VL2-23 # | | VH1-2/VL2-23 # | |
|---|---|---|---|---|---|---|---|---|
| | | | Total | mutants | Total | mutants | Total | mutants |
| 1 | 48401 | 110 | 86 | 79 | 76 | 49 | 68 | 45 |
| | 48402 | 40 | 28 | 25 | 26 | 19 | 25 | 18 |
| | 48404 | 58 | 31 | 27 | 36 | 29 | 29 | 23 |
| | 48405 | 55 | 40 | 40 | 35 | 28 | 35 | 28 |
| 2 | 48407 | 43 | 19 | 17 | 22 | 16 | 15 | 13 |
| | 48411 | 52 | 34 | 33 | 34 | 28 | 33 | 29 |
| | 48412 | 39 | 25 | 23 | 23 | 19 | 23 | 18 |
| Total | | 397 | 263 | 244 | 252 | 188 | 228 | 174 |

**Figure S6. Isolation of CH848 10.17DT-specific antibodies. Related to Figure 7.**

(**A**) Gating strategy used for single cell sort of CH848 10.17DT-specific memory B cells from vaccinated mice splenocytes.

**(B)** Summary of single B cell sort and cloning of Ig genes by PCR. A total number of 617 B cells were sorted on 96-well plates. 397 pairs of heavy and light chains were successfully recovered by sequencing of PCR amplicon. Among these heavy and light chain pairs, 263 of heavy chains were DH270 VH1-2, 252 of light chains were DH170 VL2-23, and 228 pairs were both DH270 heavy chain VH1-2 and DH270 light chain VL2-23. 169 of 228 DH270-like antibodies have acquired at least one amino acid change. DH270.mo84 was isolated from animal 48402; DH270.mo85 was isolated from animal 48411; DH270.mo86 was isolated from animal 48412.

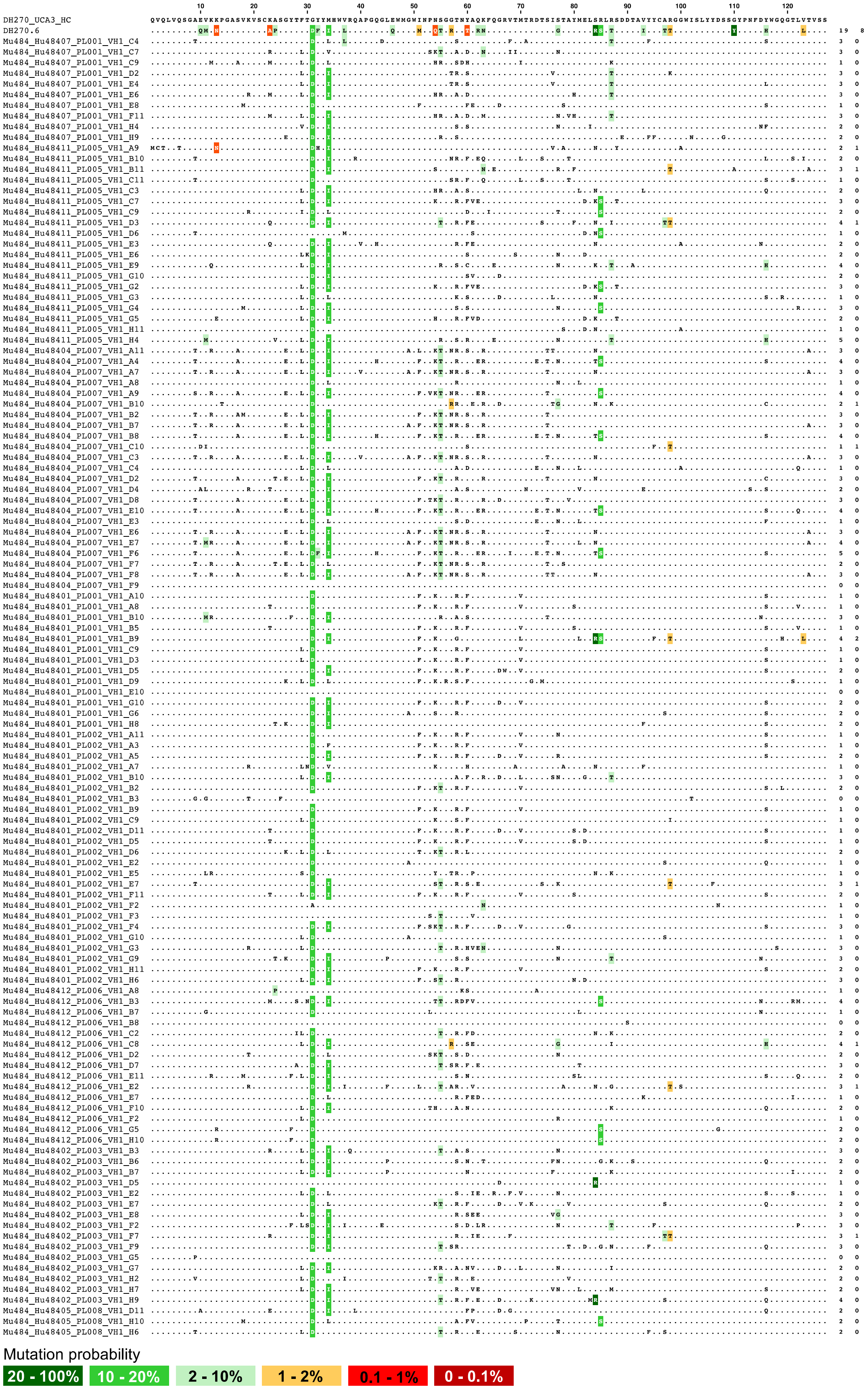
DH270_UCA3_HC
DH270.6
Mutation probability
20 - 100%
10 - 20%
2 - 10%
1 - 2%
0.1 - 1%
0 - 0.1%

**Figure S7. CH848 10.17DT trimer-ferritin NP mRNA-LNP immunization elicited antibodies with heavy chain somatic mutations shared with DH270.6. Related to Figure 7.**

Heavy chain amino acids of isolated VH1-2/VL2-23 DH270-like mAbs aligned with DH270 UCA and DH270.6. Dots represent amino acids that are in common with DH270 UCA. Mutations shared with DH270.6 are highlighted in colors according to mutation probability. The numbers to the right of each row are total numbers of probable mutations and improbable mutations observed, respectively.

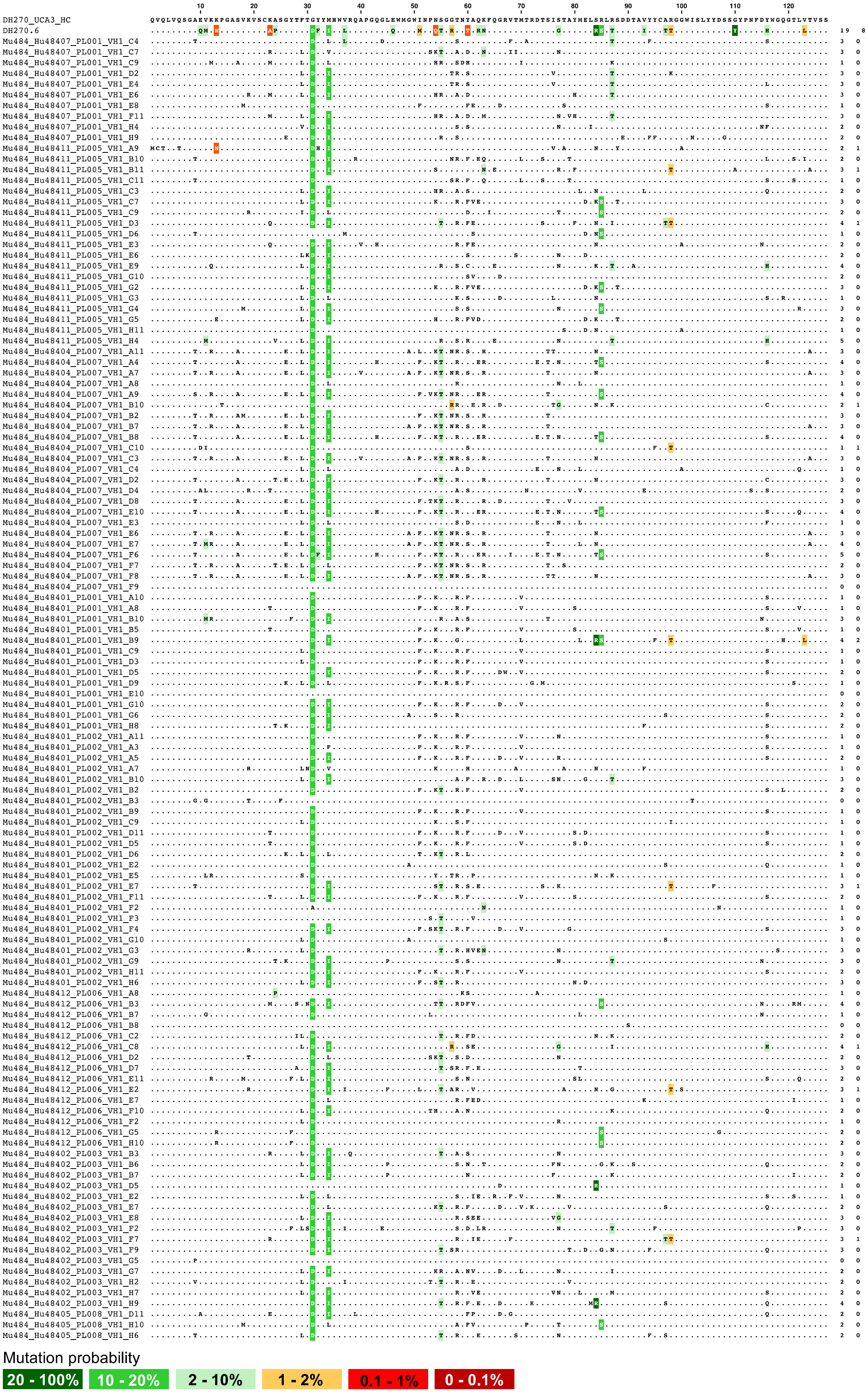
Mutation probability
20 - 100%
10 - 20%
2 - 10%
1 - 2%
0.1 - 1%
0 - 0.1%

**Figure S8. CH848 10.17DT trimer-ferritin NP mRNA-LNP immunization elicited antibodies with light chain somatic mutations shared with DH270.6. Related to Figure 7.**

Light chain amino acids of isolated VH1-2/VL2-23 DH270-like mAbs aligned with DH270 UCA and DH270.6. Dots represent amino acids that are in common with DH270 UCA. Mutations shared with DH270.6 are highlighted in colors according to mutation probability. The numbers to the right of each row are total number of probable mutations and improbable mutations, respectively.
